# Dual-trait pleiotropic analysis in highly stratified natural populations using genome-wide association summary statistics

**DOI:** 10.1101/193417

**Authors:** Xiao Feng, Yanjun Zan, Zheng Ning, Weilin Xu, Qianhui Wan, Dongyu Zeng, Ziyi Zeng, Yang Liu, Xia Shen

## Abstract

Genome-wide association analysis is a powerful tool to identify genomic loci underlying complex traits. However, the application in natural populations comes with challenges, especially power loss due to population stratification. Here, we introduce a bivariate analysis approach to a GWAS dataset of *Arabidopsis thaliana*. We demonstrate the efficiency of double-phenotype analysis to uncover hidden genetic loci masked by population structure via a series of simulations. In real data analysis, acommon allele, strongly confounded with population structure, is discovered to be associated with late flowering and slow maturation of the plant. The discovered genetic effect on flowering time is further replicated in independent datasets. Using Mendelian randomization analysis based on summary statistics from our GWAS and expression QTL scans, we predicted and replicated a candidate gene *AT1G11560* that potentially causes this association. Further analysis indicates that this locusis co-selected with flowering-time-related genes. The discovered pleiotropic genotypephenotype map provides new insights into understanding the genetic correlation of complex traits.

## Introduction

Evolution has resulted in the speciation and adaptation of various organisms. Although natural selection applies to all kinds of species, the resulting natural population structures have dramatic differences. Especially, due to their lack of mobility, plants, compared to humans and most animals, have established much stronger population structures adaptive to specific local environments (Ch. 11 in Crawley, 2009). This makes it difficult, for instance, in modern genomic studies, to distinguish genotypic effects on plants’ phenotypes from geographical stratification (Atwell et al., 2010).

Fast-developing genotyping techniques have made genome-wide association study (GWAS) one of the most useful approaches for discovering genomic loci that regulate phenotypes in various organisms(Atwell et al., 2010; Hirschhorn and Daly, 2005; Huang et al., 2010). In human GWAS, we learned that most of the discovered loci associated with complex traits or diseases have very small effects (Yang et al., 2010). The detected single nucleotide polymorphisms (SNPs) need to have sufficiently high minor allele frequencies (MAFs) for the statistical tests to gain enough power, while high-MAF variants tend to have small effects on the studied phenotypes as these variants were under weak selection pressure. Alleles that have high penetrance on a phenotype are normally under strong selection, resulting in low MAFs of the corresponding SNPs. Thus, a major challenge in human GWAS appears to be the trade-off between statistical power and the effect size of the variant to detect(Korte and Farlow, 2013; Wellenreuther and Hansson, 2016; Yang et al., 2014).

Although a similar trade-off also applies to GWAS in plant populations, e.g. in the natural population of *Arabidopsis thaliana*, in terms of discovery power, the major challenge is different. As each individual plant accession is sampled from a specific geographical location in the world, accessions with different genotypes normally have much greater phenotypic differences compared to those in humans. It appears that the genome can explain a large proportion of variation in the plant phenotype; however, the population structure in nature makes such a genomic effect heavily confounded with the environmental effect due to geographical stratification. Therefore, there can be a number of alleles, that have large genetic effects on a certain phenotype but are masked by the population structure.

As a community-based effort, over 1000 natural *A. thaliana* accessions have been collected from worldwide geographical locations (1001 Genomes Consortium, 2016; Kawakatsu et al., 2016). Most of those plants have been sequenced for genome, transcriptome, and even methylome, and these datasets have been made publicly available for worldwide researchers. Many accessions in this collection have been phenotyped for developmental, metabolic, ionomics, and stress resistance traits (Atwell et al., 2010), and more and more phenotypes are gradually releasing. Previous analyses in those datasets have revealed substantial connections between genotypic and phenotypic variations in this species. The application of association mapping has provided insights into the genetic basis of complex traits (Atwell et al., 2010; Shen et al., 2012; Wang et al., 2017), adaptation (Shen et al., 2014), and evolutionary process. Nevertheless, many essential genotype-phenotype links are still difficult to establish based on the current GWAS data, due to the substantial population stratification highly correlated with the sampling origins of the plants. Therefore, novel powerful analyses are required to further uncover the hidden genetic regulation.

Based on publicly available *A. thaliana* datasets(1001 Genomes Consortium, 2016; Atwell et al., 2010; Kawakatsu et al., 2016; Schmitz et al., 2013), here, we aim to examine the application of a bivariate analysis method that combines the discovery power of two correlated phenotypes (Shen et al., 2017), in order to map novel pleiotropic loci that simultaneously regulate both traits. We justify the ability of the summary-statistics-based approach in handling population stratification. We interpret the statistical significance with the discovered dual-trait genotype-phenotype map. We try to replicate and *in silico* functionally investigate the candidate genes that may drive such associations.

## Methods

### Genome-wide 250k SNP array genotype data and phenotype data for 199 natural *Arabidopsis thaliana* accessions

We downloaded a public dataset on the collection of 199 natural *Arabidopsis thaliana* inbred lines containing 107 phenotypes and corresponding genotypes with 214,051 SNPs available (Atwell et al., 2010). After filtering out the variants with minor allele frequencies less than 0.10, 173,220 SNPs remained.

### Whole genome re-sequencing and RNA-seq data for a population of 1,135 natural ***A. thaliana*** accessions

1,135 natural *Arabidopsis thaliana* accessions have been collected and sequenced for the whole genome and transcriptome (1001 Genomes Consortium, 2016; Kawakatsu et al., 2016). We downloaded this sequencing dataset and removed the accessions with no measured phenotype and SNPs with minor allele frequency below 0.05 and a call rate below 0.95. The final dataset includes 1001 individuals with 1,797,898 SNPs and measured flowing time at 10°C. To scan for candidate genes, we also downloaded the transcriptome dataset of a subset of this collection (*n* = 728) (Kawakatsu et al., 2016). The final eQTL scan dataset contains RNA-seq derived RPKM-values for 24,150 genes in 648 accessions whose phenotypic and genotypic data are both available.

### Whole genome re-sequencing derived SNP genotype and RNA-sequencing derived transcriptome data for a population of 144 natural ***A. thaliana*** accessions

In an earlier study, Schmitz et al. (Schmitz et al., 2013) RNA-sequenced a collection of 144 natural *A. thaliana* accessions. We downloaded this data together with their corresponding whole-genome SNP genotypes available as a part of the 1001 Genomes project (1001 Genomes Consortium, 2016; Kawakatsu et al., 2016) to replicate our SMR findings. Following the quality control procedure in(Zan et al., 2016), we removed two accessions from the data (Alst_1 and Ws_2) due to missing genotype data and two accessions (Ann_1 and Got_7) due to their low transcript call rate (16,861 and 18,693 genes with transcripts as compared to the range of 22,574 to 26,967 for the other the accessions). The final dataset used for eQTL mapping included 1,347,036 SNPs with MAF above 0.05 and call rate above 0.95 for 140 accessions, and corresponding RNA-seq derived FPKM-values for 33,554 genes.

### Single-trait genome-wide association analysis accounting for highly stratified population strucutre

For all available traits in this dataset, we first performed a mixed model-based single-trait genome-wide association analysis to generate single-trait summaries statistics. Those summary statistics were used as input for the dual-trait analysis described in the following section. To replicate our signal, we also performed a single-trait genome-wide association analysis using a collection generated in a 1001 genomes project (1001 Genomes Consortium, 2016). To correct for the population structure in these *A. thaliana* accessions, a single-trait genome-wide scan was performed based on linear mixed models, using the `polygenic` and `mmscore` procedure in GenABEL (Aulchenko et al., 2007).

### Dual-trait genome-wide association analysis in inbred lines using summary association statistics

We performed dual-trait genome scans using our recently developed multivariate analysis method implemented in the MultiABEL package (Shen et al., 2017). The method takes the whole-genome summary statistics to infer phenotypic correlation coefficients and conducts MANOVA analysis. The core test statistic of the bivariate test is Hotelling’s *T*^2^, which follows a *χ*^2^ distribution with two degrees of freedom for the two phenotypes, i.e.

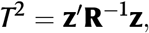

where **R** is a 2×2 matrix with the off-diagonal elements being the correlation coefficient between the two Z-scores in single-trait GWAS, for the two traits at the same variant. This correlation coefficient is proportional to the phenotypic correlation (on the liability scale for binary traits) (Zhu et al., 2015).

Thus, in the absence of population structure, the phenotypic correlation coefficient between the two traits can be unbiasedly estimated by the correlation of genome-wide Z-scores from single-trait GWAS. Linear mixed models were utilized to correct for population stratification in each single-trait GWAS; thus, intuitively, the use of the summary association statistics from single-trait analyses does not carry the issue of population stratification. However, it is unclear whether population structure affects this particular approach for estimating phenotypic correlation. Below, we first introduce the theoretical basis for this matter (see also Ning et al., 2021).

In order to model the behavior of the correlation of genome-wide association test statistics, we describe a random effect model for the phenotypic value. For *n* individuals, let the genotype vector **g** be centered, we have the phenotype vector for each trait as **y**_*i*_ = **g***β*_*i*_ + **G**_*i*_ + **e**_*i*_ (*i* = 1, 2), where *β*_*i*_ is the genetic effect at a certain variant, satisfying

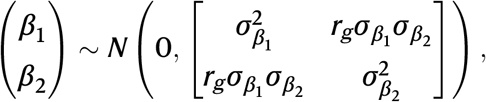

and for the rest of the genome,

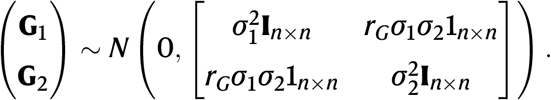

*r*_*g*_ and *r*_*G*_ are the genetic correlations between the two traits at this particular variant and the rest of the genome, respectively. Forthe *Arabidopsis* inbred lines, the phenotypes were obtained by taking the mean phenotypic value of repeated measurements for each accession (Atwell et al., 2010). This yields nearly completely genetic phenotypes, so that *e*_*i*_ approximates zero.

In single-trait GWAS, we have 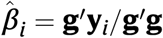, then

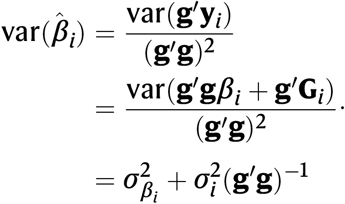

So that

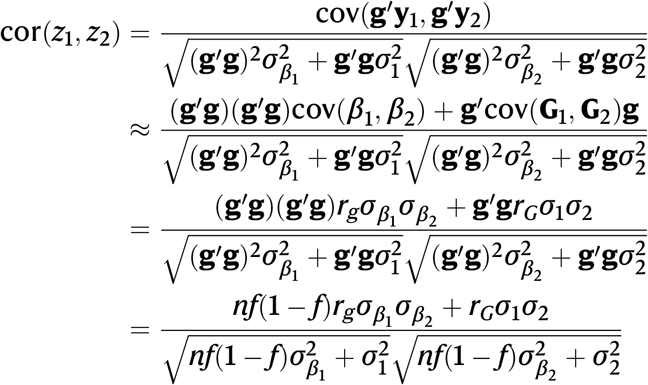

where *f* is the minor allele/genotype frequency for the inbred lines. This correlation between Z-scores serves as an estimate of phenotypic correlation because the term 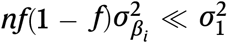, as 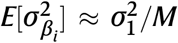 and *n ≪ M*, where *M* is the number of variants in the study. Therefore, as long as *n* is not huge, cor(*z*_1_, *z*_2_) reduces to *r*_*G*_, which is approximately the phenotypic correlation for the inbred lines population.

As the single-trait GWA analysis is performed using the linear mixed model to correct for population structure, the vector of test statistics **z** is not inflated. With the theory above, population stratification might still inflate the multivariate test statistic through genetic correlation, as the LD structure is complicatedly distributed across the genome. Neverthe less, the impact of population structure on the estimation of **R** needs to be examined empirically for each structured population.

### Simulations

In order to examine the performance of the bivariate association test in the highly stratified *Arabidopsis* population, we conducted a series of simulations. First of all, we simulated two phenotypes **y**_1_ and **y**_2_ as

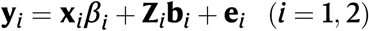

where **x**_*i*_ and **Z**_*i*_ are the genotypes for the causal SNP and the SNPs that determine population structure underlying phenotype **y**_*i*_, respectively, and *β*_*i*_ and **b**_*i*_ are the corresponding genetic effects. The population structure was simulated by the polygenic scores **Z**_*i*_**b**_*i*_ based on 10% randomly selected SNPs from the genome, excluding the SNPs within 500kb distance to the causal SNP. The polygenic effects **b**_*i*_ associated with the underlying population structure of the phenotype were drawn as **b**_*i*_ ∼ *N*(0, (*h*^2^*/M*)**I**), where *h*^2^ is the simulated narrow-sense heritability of phenotype **y**_*i*_ and *M* the number of SNPs that carry the polygenic effects. The causal SNP effect 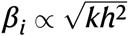, where *k* = 0, 0.001, 0.005, 0.01, 0.05. The genetic effects for the other SNPs were set to zero. The residual **e**_*i*_ ∼ *N*(0, (1 − *h*^2^)**I**). Four scenarios for the genotype data and population structure were simulated, i.e., 1) 199 natural *A. thaliana* inbred lines genotype data(173,220 SNPs) by Atwell et al.; 2) 199 natural *A. thaliana* inbred lines genotype data with half of the population structure (173,220 SNPs), where half of the individuals had their genotype data randomly permuted; 3) genotype data of the 1001 genomes project for *A. thaliana* (1,797,898 SNPs); 4) genotype data of the 1001 genomes project for *A. thaliana* with half of the population structure (1,797,898 SNPs), where half of the individuals had their genotype data randomly permuted.

The heritability value (*h*^2^) for each phenotype was set to 0.25 and 0.5 in each scenario. The phenotypic correlation (*r*_*P*_) was set to -0.25, 0, and 0.25. The simulation was repeated 1,000 times for each setup.

### eQTL and SMR analysis

We screened for candidate genes by analyzing the expression data in a subset of the 1001-genomes collection containing 140 accessions. Expression values for 19 genes around 20kb up/downstream of the top associated SNP were extracted from (Schmitz et al., 2013). Fourteen genes that did not pass the Kolmogorov-Smirnov test (KS test statistics < 0.8) were filtered out due to potential unreliable measurement mentioned in (Zan et al., 2016). The remaining five genes were subsequently passed onto eQTL mapping using qtscore procedure in GenABEL (Aulchenko et al., 2007). Outputs were reformatted according to the description in (Zhu et al., 2016). Together with the flowering time single-trait scan results (1001 Genomes Consortium, 2016), these were further passed onto SMR analysis scanning for the association between individual gene expression and flowering time. The SMR analysis was repeated for 5 top candidates, in an independent gene expression dataset containing 648 accessions (Kawakatsu et al., 2016) following the same procedure.

## Results

### Bivariate genomic scan identifies a hidden locus simultaneously associated with flowering and maturation periods

We re-analyzed a public dataset of a natural *A. thaliana* collection, where 43 developmental phenotypes and 23 flowering-time-related phenotypes were previously published (Atwell et al., 2010). The number of accessions with measured phenotypes varies from 93 to 193, with a median of 147 (Table S1). We first excluded all variants with minor allele frequencies (MAF) less than 0.1 and performed single-trait GWA analysis for all these traits based on a linear mixed model, so that the confounded genetic effects due to population stratification are adjusted. We then applied our recently developed multi-trait GWAS method (Shen et al., 2017) to all pairwise combinations of the phenotypes (see Methods). One novel locus, in one of the pairwise tests, reached the most stringent 5% Bonferroni-corrected genomewide significance threshold for the 2,145 pairs of traits and 173,220 variants, i.e. *p* < 1.35 × 10^−10^ (**Table 1, Figure 1a**). This signal also reaches single-trait genome-wide significance in the other six pairs of traits highly correlated with the top pair (Figure S1), without Bonferroni correction for the number of tested trait pairs (**Table 1**, Figure S2-S7).

**Table 1:** Discovery and replication analyses results for the novel pleiotropic locus. Reported association statistics are for the top variant at the locus for each pair of traits. ^1^LD: Days to flowering time under Long Day. ^2^0W: Days to flowering time under long day without vernalization. ^3^2W: Days to flowering time under long day with vernalized for 2 weeks at 5°C, 8hrs daylight. ^4^4W: Days to flowering time under long day with vernalized for 4 weeks at 5°C, 8 hrs daylight. ^5^0W GH FT: Days to flowering time (greenhouse). ^6^FT GH: Days to flowering (greenhouse). ^7^MT GH: Maturation period (greenhouse), 20°C, 16 hrs daylight. ^8^RP GH: Reproduction period (greenhouse), 20°C, 16 hrs daylight. ^9^RA: Reference allele. ^10^EA: Effect allele. ^11^MAF: Minor allele frequency. ^12^Correlation refers to observed phenotypic correlation. ^13^FT: Flowering time.

| *dual-trait Analysis* |  |  |  |  |  |  |  |  |  |
| --- | --- | --- | --- | --- | --- | --- | --- | --- | --- |
| Trait 1 | Trait 2 | Chr | Position | RA<sup>9</sup> | EA<sup>10</sup> | MAF<sup>11</sup> | *P* | Correlation<sup>12</sup> |  |
| LD<sup>1</sup> | MT GH<sup>7</sup> | 1 | 3895353 | C | T | 0.20 | 6.3×10<sup>−9</sup> | -0.39 |  |
| OW<sup>2</sup> | MT GH<sup>7</sup> | 1 | 3896072 | G | T | 0.20 | 8.4×10<sup>−9</sup> | -0.58 |  |
| 2W<sup>3</sup> | MT GH<sup>7</sup> | 1 | 3906923 | T | C | 0.22 | 9.9×10<sup>−12</sup> | -0.68 |  |
| 2W<sup>3</sup> | RP GH<sup>8</sup> | 1 | 3978064 | A | C | 0.27 | 1.3×10<sup>−8</sup> | -0.17 |  |
| 4W<sup>4</sup> | MT GH<sup>7</sup> | 1 | 3906923 | T | C | 0.22 | 3.1×10<sup>−9</sup> | -0.64 |  |
| OW GH FT<sup>5</sup> | MT GH<sup>7</sup> | 1 | 3906923 | T | C | 0.22 | 1.8×10<sup>−8</sup> | -0.36 |  |
| FT GH<sup>6</sup> | MT GH<sup>7</sup> | 1 | 3896072 | G | T | 0.20 | 1.5×10<sup>−8</sup> | -0.60 |  |
| *Single-trait Analysis* |  |  |  |  |  | *Replication* |  |  |  |
| Trait 1 |  |  | Trait 2 |  |  | FT<sup>13</sup> 10°C |  | FT<sup>13</sup> 16°C |  |
| Effect | *P* | *h*<sup>2</sup> | Effect | *P* | *h*<sup>2</sup> | Effect | *P* | Effect | *P* |
| 33.5 | 5.6×10<sup>−6</sup> | 0.22 | 2.42 | 6.0×10<sup>−4</sup> | 0.07 | 2.40 | 9.99×10<sup>−3</sup> | 4.52 | 8.60×10<sup>−4</sup> |
| 17.3 | 1.6×10<sup>−4</sup> | 0.17 | 2.59 | 2.1×10<sup>−4</sup> | 0.09 | 2.40 | 9.99×10<sup>−3</sup> | 4.52 | 8.60×10<sup>−4</sup> |
| 15.3 | 2.3×10<sup>−5</sup> | 0.24 | 2.47 | 3.7×10<sup>−5</sup> | 0.10 | 2.52 | 5.65×10<sup>−3</sup> | 4.44 | 7.87×10<sup>−4</sup> |
| 19.7 | 6.8×10<sup>−7</sup> | 0.26 | 2.65 | 1.6×10<sup>−3</sup> | 0.06 | 1.87 | 3.60×10<sup>−2</sup> | 2.82 | 3.22×10<sup>−2</sup> |
| 11.6 | 1.7×10<sup>−3</sup> | 0.16 | 2.47 | 3.7×10<sup>−5</sup> | 0.10 | 2.52 | 5.65×10<sup>−3</sup> | 4.44 | 7.87×10<sup>−4</sup> |
| 25.8 | 3.8×10<sup>−5</sup> | 0.21 | 2.47 | 3.7×10<sup>−5</sup> | 0.10 | 2.52 | 5.65×10<sup>−3</sup> | 4.44 | 7.87×10<sup>−4</sup> |
| 14.9 | 1.8×10<sup>−3</sup> | 0.11 | 2.59 | 2.1×10<sup>−4</sup> | 0.09 | 2.40 | 9.99×10<sup>−3</sup> | 4.52 | 8.60×10<sup>−4</sup> |

**Figure 1:**
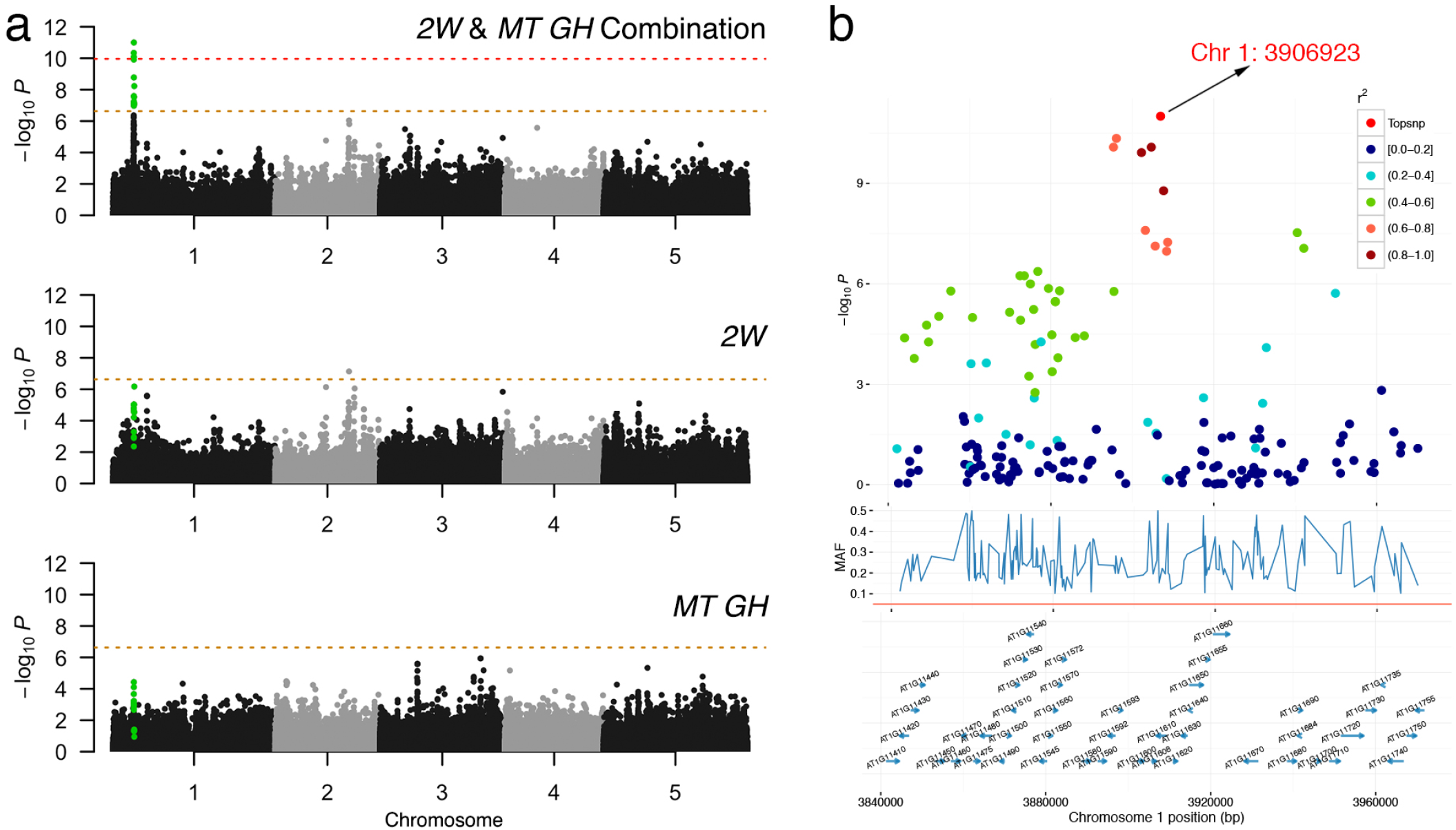
Bivariategenome-wide association analysis of two developmental traits. 2W: Days to flowering time (FT) under long day (LD) with vernalized for 2 wks at 5°C, 8hrs daylight, MT GH: Maturation period. a) Manhattan plots comparison of bivariate and univariate analysis results, where the novel variants only discoverable when combining two phenotypes are shown in green. Thehorizontal dashed line represents a 5% Bonferroni-corrected genome-wide significant threshold for the number of variants and also the number of tested trait pairs, respectively. b) Zooming in on the novel locus detected using bivariate analysis. r: linkage disequilibrium measured as the correlation coefficient between the top variant and each variant in the region.

For the most significant trait combination, 2W (days to flowering time under a long day with vernalized for 2 weeks) and MT GH (maturation period), the linkage disequilibrium (LD) block of this locus (LD *r* > 0.7) covers about a 260 kb interval on chromosome 1, with a top variant at 3,906,923 bp (dual-trait *p* = 9.9 × 10^−12^, **Figure 1b, Table 1**). The detected locus shows joint effects on flowering and maturation, where the effect on flowering time (2W) is notably large (15.3 days), and that on maturation period (MT GH) is 2.5 days (**Table 1**). These correspond to narrow-sense heritability values of 24% and 10% of the two phenotypes, respectively.

### dual-trait analysis is sufficiently powerful to overcome the confounding population structure

The detected joint-effect locus was missed in the corresponding single-trait GWA analysis of 2W (effect = 15.3, *p* = 2.26 × 10^−5^ after correcting for population stratification) and that of MT GH (effect = 2.5, *p* = 3.70 × 10^−5^). Notably, this locus was not even detectable at the genome-wide significance level in a much larger population of more than 1,000 *A. thaliana* accessions (1001 Genomes Consortium, 2016; Kawakatsu et al., 2016) due to its severe confounding with the natural population structure. The statistical significance can only be identified when considering the joint distribution of the bivariate statistic. According to the genome-wide Z-scores (student t-statistics), these two phenotypes are negatively correlated, as the plant’s lifespan is relatively stable (estimated and observed phenotypic correlation = -0.55 and -0.68, respectively). However, the observed effects on the two traits are both substantially positive, showing sufficient deviation from the joint distribution that led to bivariate statistical significance (**Figure 2**).

**Figure 2:**
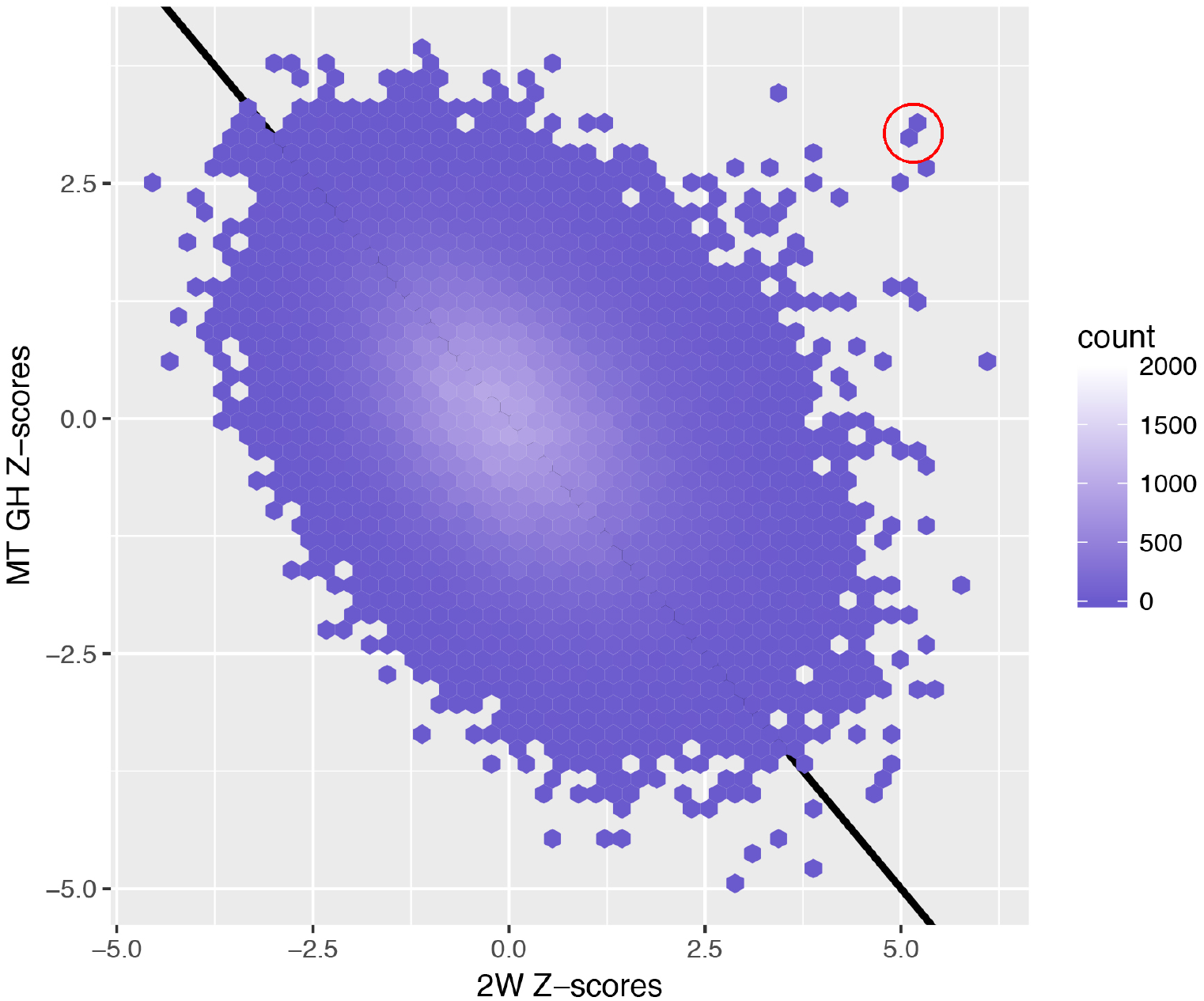
Hexbin scatter plot comparing all Z-scores of the two traits across the genome, showing the bivariate statistical significance of the detected locus. The top variants of the locus are marked on the edge of the empirical bivariate normal distribution with a red circle. The black line with a slope of -1 is provided as a visual guide.

The strong confounding with the population structure can also be visualized by the allele frequency distribution of the top associated SNP across different *A. thaliana* subpopulations based on the genome re-sequencing data from the *A. thaliana* 1001-genomes project (1001 Genomes Consortium, 2016) (**Figure 3**). The sub-populations were divided by admixture analysis using ADMIXTURE (1001 Genomes Consortium, 2016; Alexander et al., 2009). The plus allele increasing flowering time was predominantly found in Sweden and almost fixed in the Northern Sweden population(**Figure 3b**; allele frequency= 0.97 in Northern Sweden and 0.51 in Southern Sweden). Overall, the phenotype, e.g. flowering time at 10 °C, highly correlates with the frequency of the plus allele (**Figure 3**). The genotype at this locus follows a latitude decline, where the northern accessions are enriched with the plus allele, and the southern accessions are enriched with the minus allele (**Figure 3**). This spatially imbalanced enrichment shows strong confounding with the population structure, which is why standard single-trait GWAS loses power substantially.

**Figure 3:**
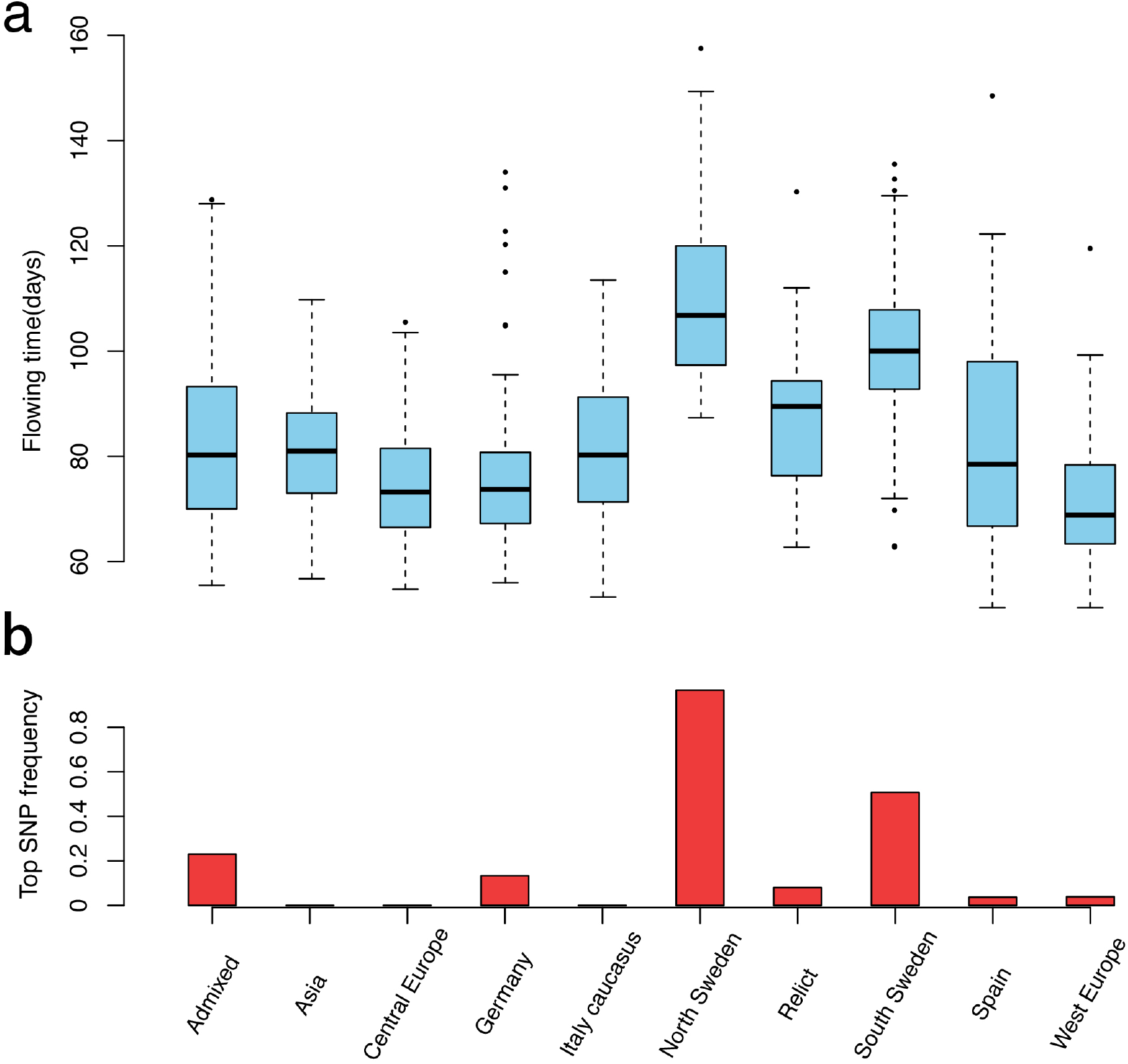
The discovered locus is highly confounded with population structure. a) Flowering time variation (10°C) among different sub-populations of *Arabidopsisthaliana*. These populations are divided by admixture analysis (1001 Genomes Consortium, 2016); b) Frequency of the top associated SNP at chromosome 1, 3,906,923 bp in different subpopulations. The association between the structure of the phenotype and that of the allele frequency shows the population confounding at this locus.

### Summary-level dual-trait analysis controls confounding-induced statistical bias

The analysis procedure above is intuitively straightforward, without sophisticated bivariate linear mixed modeling. Although the linear mixed model was incorporated in the genomewide association scan for every single phenotype, it is unclear whether the dual-trait analysis using single-trait summary association statistics can control the confounding due to the substantial structure in the *A. thaliana* natural population, especially given that longrange LD is common in such populations (Long et al., 2013).

In order to justify our summary-statistics-based approach, we performed a series of simulations (see Methods). The simulation scenarios cover different levels of population stratification, levels of heritability for two phenotypes, various levels of the effect of the causal SNP on each phenotype, and the situations where the directions of causal effects on the two phenotypes are consistent or inconsistent with the phenotypic correlations (Figure S8-S11). First of all, we found that the summary-statistics-based dual-trait test was able to control the false positive rates, given that the summary statistics were from the single-trait linear mixed model (**Figure 4**, column 1; Figure S12). As expected, if the causal SNP only has an effect on one of the two phenotypes, the dual-trait test would have a slightly lower power that the single-trait analysis (**Figure 4**, column 4). Nevertheless, when the SNP has pleiotropic effects on both phenotypes, the dual-trait analysis has better power in detecting the genetic association, especially when the genetic and phenotypic correlations at the causal SNP have opposite directions(**Figure 4**, column 5).

**Figure 4:**
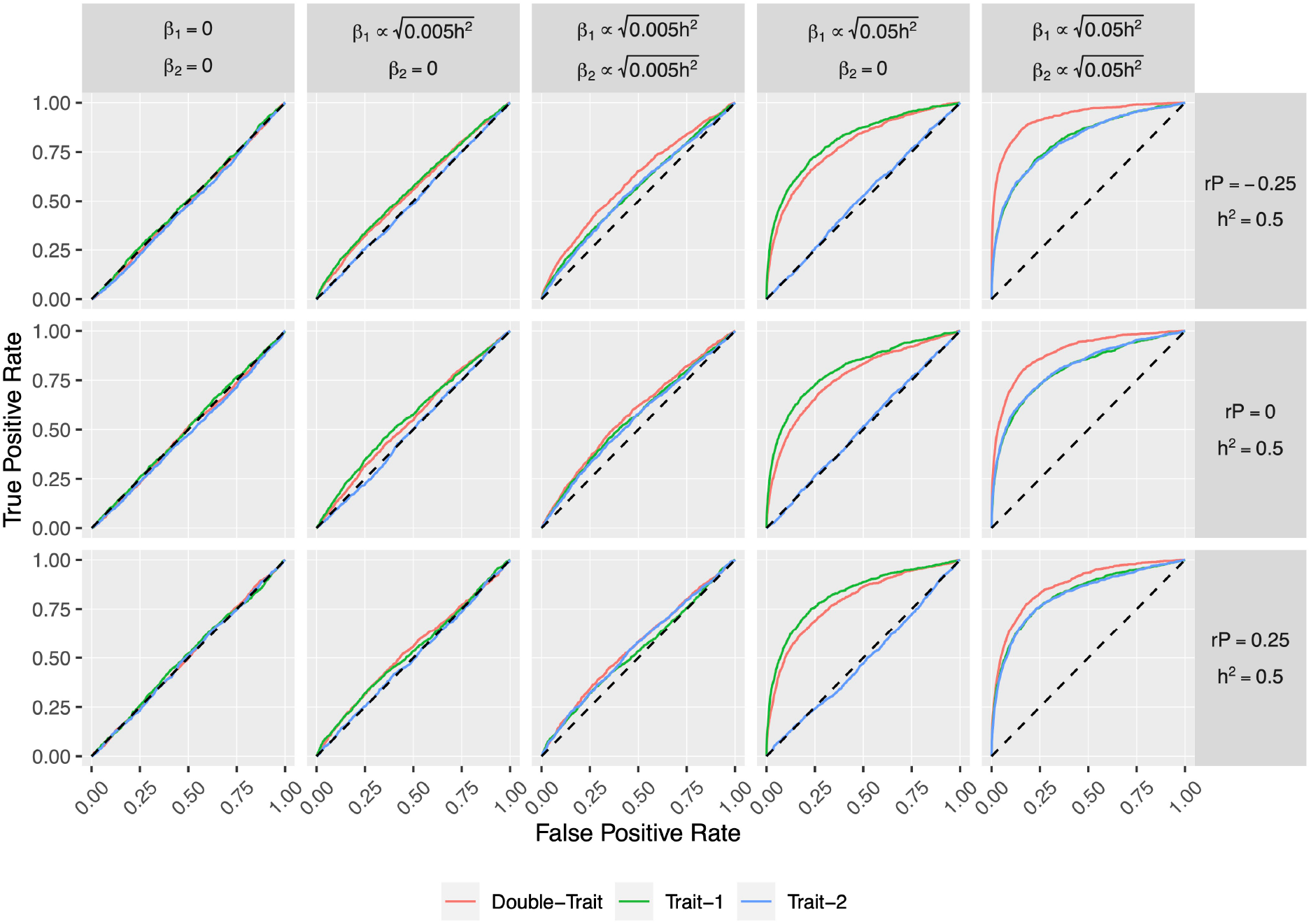
Comparisons of single-trait and dual-trait GWAS results of the target SNP under different simulated scenarios using 199 natural *Arabidopsisthaliana* inbred lines genotype data. For each scenario, ROC curves represent the true positive rate against the false positive rate with single- and dual-trait GWAS methods from 1000 simulations. The target SNP’s effect size (*β*) of each phenotype was indicated at the top of ROC plots. The heritability (*h*^2^) of each phenotype was set to 0.5, with 10% genome-wide SNPs randomly chosen to simulate population structure. The phenotypic correlation (*r*_*P*_) was set to -0.25, 0, and 0.25, respectively. More scenarios can be found in Figures. S8-11.

### Replication of the detected genetic effect on flowering time

Although we lack an independent dataset of *A. thaliana* maturation duration to replicate the bivariate statistical test, datasets containing additional independent *A. thaliana* flowering time measurements are available. We downloaded a flowering time GWAS dataset Measured in 1,135 natural accessions from the 1001-genomesprojectcollection(1001 Genomes Consortium, 2016) and performed a single-trait association analysis of our discovered top SNP with linear mixed model correction for the population structure. The genetic effect was significantly replicated for flowering time at 10 °C (effect = 2.5 days, *p* = 0.006) and flowering time at 16 °C (effect = 4.4 days, *p* = 0.0008). The effects on flowering time in the replication sample appear to be smaller than in the discovery population, possibly due to Winner’s curse in the discovery phase.

We also screened literature for conventional quantitative trait loci (QTL) studies in intercrosses using natural *A. thaliana* accessions. Our detected signal is underneath a reported QTL peak for flowering time from an intercross between a Swedish and an Italian accession (Dittmar et al., 2014) (Figure S13). This, together with the replication above, justifies the detected association. Although the discovered genetic effect on the maturation period is not directly replicated, the effect does exist when the effect on flowering is justified, as the pleiotropic signal must be driven by both phenotypes.

### Prediction and replication of candidate genes using summary-level Mendelian randomization

As a community-based effort, all the natural *A. thaliana* accessions from the 1001-genomes project were measured for their transcriptome(1001 GenomesConsortium, 2016; Kawakatsu et al., 2016). Such a public gene expression dataset allows us to predict candidate genes underlying the association signal. We extracted the expression levels of 19 genes within a *±* 20kb window around the top associated SNP using RNA-seq gene expression measurements from 140 accessions (Schmitz et al., 2013). Among these, the distributions of 14 gene expression phenotypes significantly deviate from normality (Kolmogorov-Smirnov test statistic > 0.8), and these genes were filtered out due to potentially unreliable measurements (Zan et al., 2016). The remaining 5 genes were passed onto eQTL mapping at the discovered locus (see Methods).

Based on the locus-specific eQTL mapping summary statistics, we applied the recently developed Summary-level Mendelian Randomization (SMR) method (Zhu et al., 2016) to predict potential candidate genes among these five genes. The analysis integrates summary association statistics from GWAS and eQTL scan to predict functional candidate genes using multiple-instrument Mendelian randomization (Burgess et al., 2015), where the complementary HEterogeneity In Dependent Instruments(HEIDI) test checks that the gene expression and flowering time share the same underlying causal variant. One significant can-didate *AT1G11560* was detected after Bonferroni correction for five tests (**Figure 5, Table 2**). This candidate gene prediction result was also replicated using an independent eQTL mapping dataset (Kawakatsu et al., 2016).

**Figure 5:**
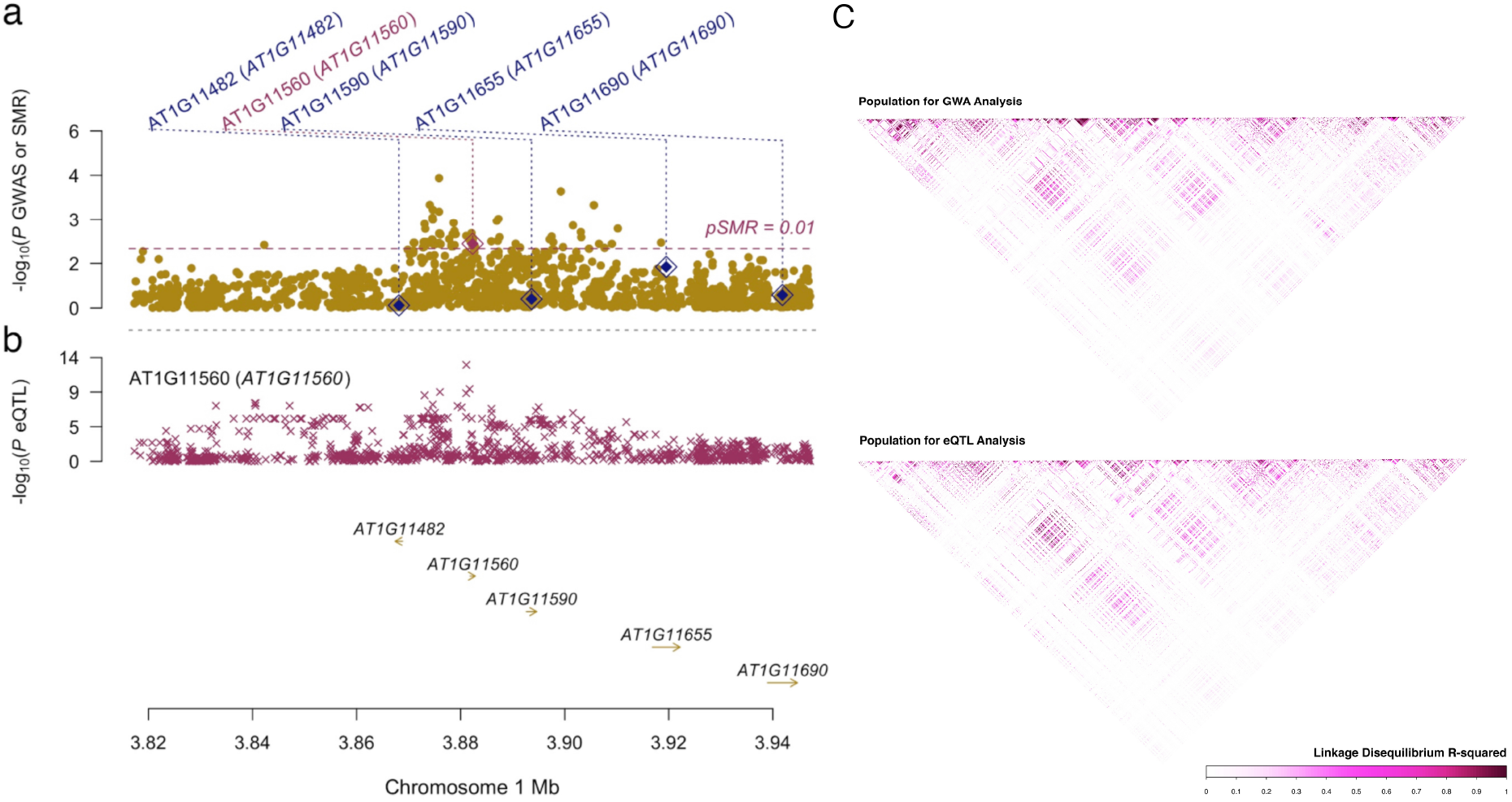
Prioritized candidate genes at the detected locus for flowering time using SMR analysis. a) Manhattan plot of the association between the flowering time at 10°C and SNPs around 40kbof the top associated SNP in bivariate analysis. The diamonds highlight top eQTL for individual genes. b) Manhattan plot of the association between expression of *AT1G11560* and SNPs around 40kb of the top associated SNP in bivariate analysis. Genes tested in SMR analysis are highlighted using arrows. c) Similar linkage-disequilibrium structure at the locus for the corresponding populations of GWA and eQTL analyses.

**Table 2:**
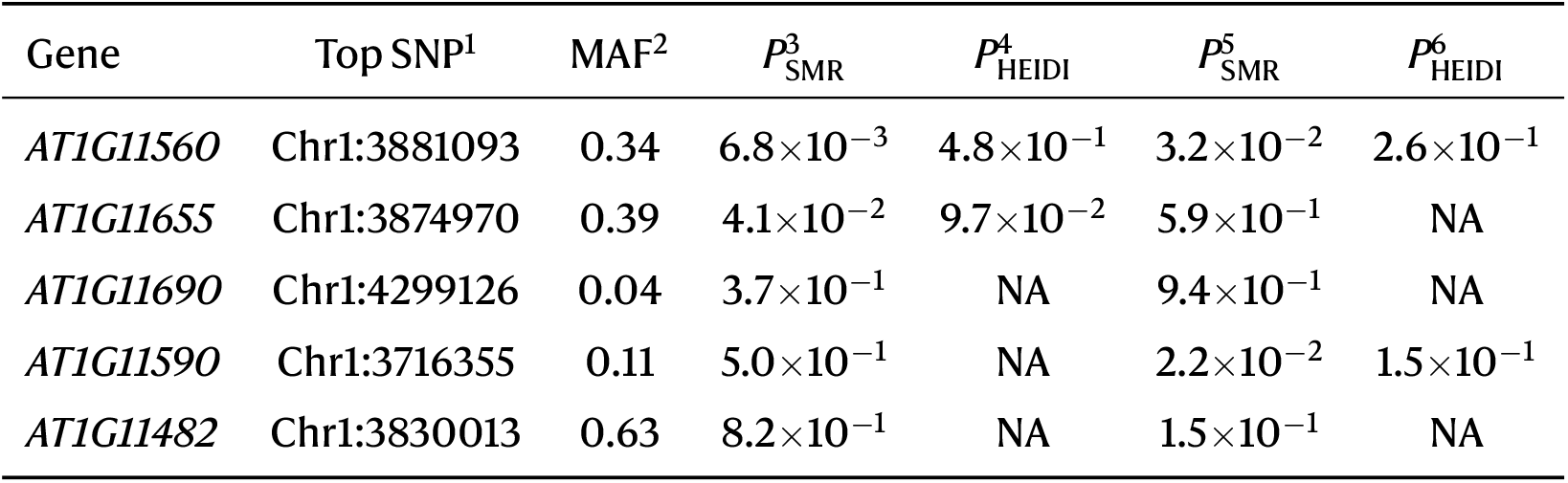
Summary of the SMR/HEIDI analysis results. ^1^Top SNP: The top SNP in expression QTL analysis. ^2^MAF: Minor allele frequency of the top associated SNP. ^3^*P*_SMR_: p-value from SMR using a collection of 140 *A. thaliana* accessions. ^4^*P*_HEIDI_: p-value from HEIDI test using a collection of 140 *A. thaliana*. ^5^*P*_SMR_: p-value from SMR using a second collection of 648 accessions. ^6^*P*_HEIDI_: p-value from HEIDI test using a second collection of 648 accessions.

### Indication of co-selection with genes in flowering-related pathways

As flowering time is a well-known polygenic trait, we expect multiple loci to be involved and possibly co-selected as a result of parallel evolution. Therefore, we explored the evidence of co-selection by associating the expression values of 288 known genes in floweringtime-related pathways and one gene in the maturation-related pathway with our top SNP using transcriptome data from 648 *A. thaliana* accessions (1001 Genomes Consortium, 2016) (see Methods). In total, six genes (*NF-YA8, AT5G53360, SPL15, AGL42, FLC, AGL20*) were associated with our top SNP (false discovery rate < 0.05), where, conservatively, four genes (*AT5G53360, AGL42, FLC, AGL20*) were replicated after Bonferroni correction for six tests using data from an independent collection of 140 *A. thaliana* (Schmitz et al., 2013) (Table S2). This indicates that co-selected genes in multiple pathways determine the flowering time variation in nature, and our detected locus contributes to a part of that.

## Discussion

A serious issue of GWAS in natural population is the confounding between true underlying genetic effects and the population structure, which can lead to spurious associations between genotypes and phenotypes if population stratification is not properly adjusted (Korte and Farlow, 2013; Wellenreuther and Hansson, 2016; Yang et al., 2014). In corporation of the random polygenic effect using linear mixed models can effectively control the population structure, but such correction often compromises the true signals. Here, we applied a bivariate analysis to a classic dataset and successfully separated a locus from the strong population structure. The detected allele is associated with late flowering and slow maturation of *A. thaliana*, which was corrected away by the linear mixed model in a standard single-trait analysis. The replication of the genetic effect on flowering time in an old intercross linkage analysis and another independent dataset improves the confi-Dence of this association. The discovered association is atypical example that jointly modeling phenotypes that share the genetic basis can boost discovery power and reveal the pleiotropic genotype-phenotype map at the same time.

Together with our recent application of multivariate analysis in human isolated populations (Shen et al., 2017), the results further indicate that multi-phenotype analysis is an effective approach to detect hidden loci that lack discovery power in single-phenotype analysis and thus is worth testing in broader applications. The multivariate analysis appears to have the greatest power when the locus-specific genetic correlation does not agree with the natural phenotypic correlation. For instance, like the discovery here, for two traits that are negatively correlated, loci that generate a positive genetic correlation between the traits tend to have a good chance of being detected in a joint analysis.

In GWAS, phenotypes are usually chosen based on morphological, physiological, or economical features. Those features are usually feasible and simple to quantify; however, they might not be directly representative of the underlying genetic or biological factor that we try to detect. Fortunately, a certain degree of biological pathway sharing among complex traits is common, i.e. pleiotropy (Visscher and Yang, 2016). Nowadays, it is very common that multiple phenotypes are measured for the same individuals in many GWAS datasets, especially in omics studies where thousands of phenotypes are measured. Instead of focusing on one phenotype at a time, it is of essential value to jointly model multiple phenotypes, attempting to detect pleiotropic loci that affect multiple traits with biological relevance.

In this study, all the pairs of traits that are associated with the detected locus contain at least one flowering-time trait, and nearly all of them have maturation duration involved. Detection of the novel locus in a bivariate analysis indicates a shared genetic basis for the two types of developmental traits, which measure the lengths of two important periods during the plant’s lifetime. By integrating the expression level information and GWAS result using SMR/HEIDI test, we were able to predict candidate genes in this region. However, further work beyond the scope of this paper is still required to establish the molecular biological basis underlying the replicate association.

Many genetic variants affecting flowering time have been mapped, and many genes promoting flowering times have been well characterized using standard lab accession, Col-0 (Brachi et al., 2010). Unlike simple traits, where only one or a few alleles are driving the trait’s variation, there are many more variants throughout the genome that contribute to the variation of flowering time. The associations between our top SNP and the expression of many flowering-time-related genes serve as evidence of co-selection or parallel adaptation.

Inconclusion, our study demonstrates the efficiency of joint modeling multiple phenotypes which overcomes severe power loss due to population stratification in association studies. We discover and replicate a pleiotropic allele that regulates flowering and maturation periods simultaneously, providing novel insights into understanding the plant’s development over a lifetime. By integrating gene expression information with the GWAS results, we predict a functional candidate underneath the associated genomic region. We encourage wider applications of such a multivariate framework in future analyses of genomic data.

## Acknowledgements

X.S. was in receipt of a National Natural Science Foundation of China (NSFC) grant (No. 12171495), a Natural Science Foundation of Guangdong Provincegrant(No. 2114050001435), a National Key Researchand Development Programgrant(No. 2022YFF1202105), and Swedish Research Council (Vetenskapsrådet) grants (No. 2014-00371, No. 2017-02543, No. 2022-01309). International collaboration within this work was partly supported by the Swedish Foundation for International Cooperation in Research and Higher Education (STINT) initiation grant to X.S. (No. IB2015-6000) and Karolinska Institutet travel grant (No. 2017-00534). The funders had no role in study design, data collection and analysis, the decision to publish, or the preparation of the manuscript.

## Author contributions

X.S. initiated and coordinated the study. X.S. and Y.L. supervised the study. X.F. and Y.Z. performed the data analysis. Z.N. and X.S. contributed to statistical modeling and interpretation. W.X., Q.W., D.Z. and Z.Z. contributed to data processing. X.F., Y.Z. and X.S. wrote the manuscript. All authors approved the final version of the manuscript.

## Data availability

All genotypes and phenotypes data we used are publicly available from 1001GenomesConsortium (2016); Atwell et al. (2010); Kawakatsu et al. (2016); Schmitz et al. (2013); Zan et al. (2016). Atwell et al.’s 199 natural *Arabidopsis thaliana* inbred lines containing 107 phenotypesand correspondinggenotypes are publicly available at https://github.com/Gregor-Mendel-Instituatpolydb/blob/master/miscellaneous_data/phenotype_published_raw.tsv and https://github.com/Gregor-Mendel-Institute/atpolydb/blob/master/250k_snp_data/call_method_75.tar.gz.

