## Supplementary Information for "Dual-trait pleiotropic analysis in highly stratified natural populations using genome-wide association summary statistics"

#### Contents

**1 Supplementary Tables** **2**

**2 Supplementary Figures** **5**

### 1 Supplementary Tables

**Table S1: Phenotypes included in the bivariate analyses.** Details about phenotyping can be referred to Atwell et al. (2010).

| Phenotype | Description | Number of Accessions |
|---|---|---|
| LD | Days to flowering time (FT) under Long Day (LD) | 167 |
| LDV | Days to flowering time (FT) under Long Day (LD) (5 wks vernalization) | 168 |
| SD | Days to flowering time (FT) under Short Day (SD) | 162 |
| SDV | Days to flowering time (FT) under Short Day (SD) (5 wks vernalization) | 159 |
| 0W | Days to FT under LD without vernalization | 137 |
| 2W | Days to FT under LD with 2wks vernalization | 152 |
| 4W | Days to FT under LD with 4wks vernalization | 119 |
| 8W | Days to FT under LD with 8wks vernalization | 155 |
| FLC | FLC gene expression | 167 |
| FRI | FRI gene expression | 164 |
| FT10 | Flowering time (FT), 10°C | 194 |
| FT16 | Flowering time (FT), 16°C | 193 |
| FT22 | Flowering time (FT), 22°C | 193 |
| LN10 | leaf number at flowering time (LN), 10°C | 177 |
| LN16 | leaf number at flowering time (LN), 16°C | 176 |
| LN22 | leaf number at flowering time (LN), 22°C | 176 |
| 8W GH FT | Days to FT with 8 wks vernalization | 162 |
| 8W GH LN | LN at FT with 8 wks vernalization | 163 |
| 0W GH FT | Days to FT without vernalization | 153 |
| 0W GH LN | LN at FT without vernalization | 135 |
| FT Field | Days to flowering of plants grown in the field | 180 |
| FT Diameter Field | Plant diameter at flowering (field) | 180 |
| FT GH | Days to flowering (greenhouse) | 166 |
| LES | Presence or absence of lesioning | 95 |
| YEL | Presence or absence of yellowing | 95 |
| LY | Presence or absence of either lesioning or yellowing | 95 |
| FW | Fresh weight of plants | 95 |
| DW | Dry weight of plants | 95 |
| Chlorosis 10 | Visual chlorosis presence, 10°C | 177 |
| Chlorosis 16 | Visual chlorosis presence, 16°C | 176 |
| Chlorosis 22 | Visual chlorosis presence, 22°C | 176 |
| Anthocyanin 10 | Visual anthocyanin presence, 10°C | 177 |
| Anthocyanin 16 | Visual anthocyanin presence, 16°C | 176 |
| Anthocyanin 22 | Visual anthocyanin presence, 22°C | 177 |
| Seed Dormancy | Seed dormancy level | 83 |
| Germ 10 | Days to germination, 10°C | 177 |
| Germ 16 | Days to germination, 16°C | 176 |
| Germ 22 | Days to germination, 22°C | 177 |
| Seedling Growth | Seedling growth rate | 100 |
| Vern Growth | Vegetative growth rate during vernalization | 110 |
| After Vern Growth | Vegetative growth rate after vernalization | 110 |
| Secondary Dormancy | Decrease in germination rate after prolonged exposure to cold temperature | 93 |
| Germ in dark | Germination in the dark | 93 |
| DSDS50 | Duration of seed dry storage required for 50\% of the seeds to germinate | 109 |
| Seed bank 133-91 | Non-monotonous dynamic of dormancy release | 110 |
| Storage 7 days | Primary dormancy, 7 days dry storage | 110 |
| Storage 28 days | Primary dormancy, 28 days dry storage | 110 |

| | | |
|---|---|---|
| Storage 56 days | Primary dormancy, 56 days dry storage | 110 |
| Hypocotyl length | Hypocotyl length | 89 |
| Width 10 | Plant diameter, 10°C | 176 |
| Width 16 | Plant diameter, 16°C | 175 |
| Width 22 | Plant diameter, 22°C | 175 |
| Leaf serr 10 | Level of leaf serration, 10°C | 174 |
| Leaf serr 16 | Level of leaf serration, 16°C | 176 |
| Leaf serr 22 | Level of leaf serration, 22°C | 176 |
| Leaf roll 10 | Leaf roll presence, 10°C | 177 |
| Leaf roll 16 | Leaf roll presence, 16°C | 176 |
| Leaf roll 22 | Leaf roll presence, 22°C | 176 |
| Rosette Erect 22 | Presence of rosette errectness, 22°C | 176 |
| Silique 16 | Silique length, 16°C | 95 |
| Silique 22 | Silique length, 22°C | 95 |
| FT Duration GH | Flowering period duration | 147 |
| LC Duration GH | Life cycle period | 147 |
| LFS GH | Last flower senescence | 148 |
| MT GH | Maturation period | 147 |
| RP GH | Reproduction period | 147 |

**Table S2: Genes in flowering-time pathways whose expression are associated with the detected locus.** [1]p-value from a expression dataset generated from 648 accessions in the *A. thaliana* 1001-genomes project (Kawakatsu et al., 2016). [2]FDR value computed from p-value. [3]Replication p-value from another subset of 140 accessions (Schmitz et al., 2013).

| Locus ID | Gene Name | p-value[1] | q-value[2] | Replication p-value[3] |
|---|---|---|---|---|
| AT1G17590 | *NF-YA8* | $1.6\times10^{-7}$ | $2.3\times10^{-5}$ | $1.7\times10^{-2}$ |
| AT5G53360 | *AT5G53360* | $5.8\times10^{-7}$ | $5.7\times10^{-5}$ | $3.2\times10^{-4}$ |
| AT3G57920 | *SPL15* | $7.9\times10^{-4}$ | $7.8\times10^{-3}$ | $1.7\times10^{-2}$ |
| AT5G62165 | *AGL42* | $1.2\times10^{-3}$ | $1.1\times10^{-2}$ | $6.3\times10^{-3}$ |
| AT5G10140 | *FLC* | $1.5\times10^{-3}$ | $1.3\times10^{-2}$ | $5.7\times10^{-4}$ |
| AT2G45660 | *AGL20* | $1.8\times10^{-3}$ | $1.4\times10^{-2}$ | $1.2\times10^{-3}$ |

#### 2 Supplementary Figures

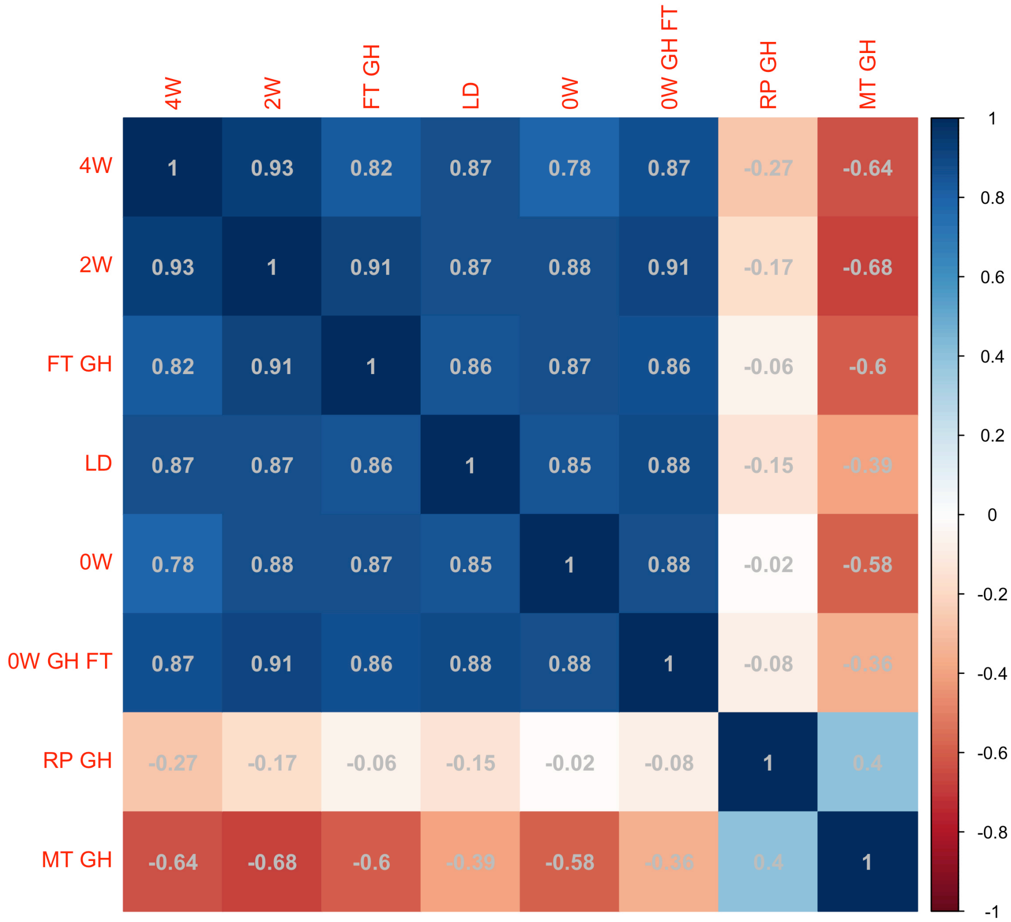


**Figure S1: Phenotypic correlations among flowering time-related traits, reproduction period, and maturation period phenotypes.** The flowering time-related traits are: 4W: Days to flowering time (FT) under long day (LD) with vernalized for 4 wks at 5°C, 8hrs daylight; 2W: Days to flowering time (FT) under long day (LD) with vernalized for 2 wks at 5°C, 8hrs daylight; FT GH: Days to flowering (greenhouse); LD: Days to flowering time (FT) under Long Day (LD); 0W: Days to flowering time (FT) under Long Day (LD) without vernalization; 0W GH FT: Days to flowering time (FT) without vernalization.

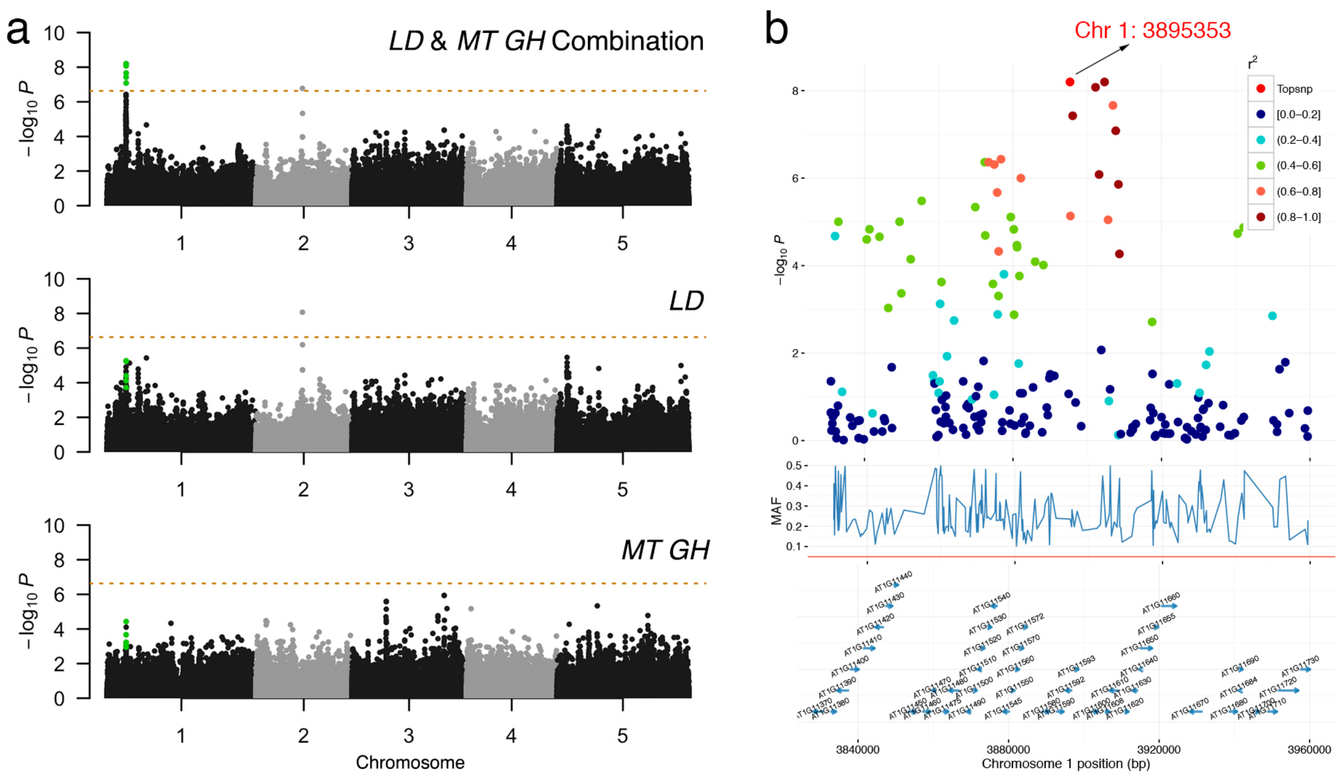


**Figure S2: Bivariate genome-wide association analysis of two developmental traits, LD: Days to flowering time (FT) under Long Day (LD), MT GH: Maturation period.** (a) Manhattan plots comparison of bivariate and univariate analysis results, where the novel variants are only discoverable when combining two phenotypes are shown in green. The horizontal dashed line represents a 5% Bonferroni-corrected genome-wide significant threshold. (b) Zooming in on the novel locus detected using bivariate analysis. $r$: linkage disequilibrium measured as the correlation coefficient between the top variant and each variant in the region.

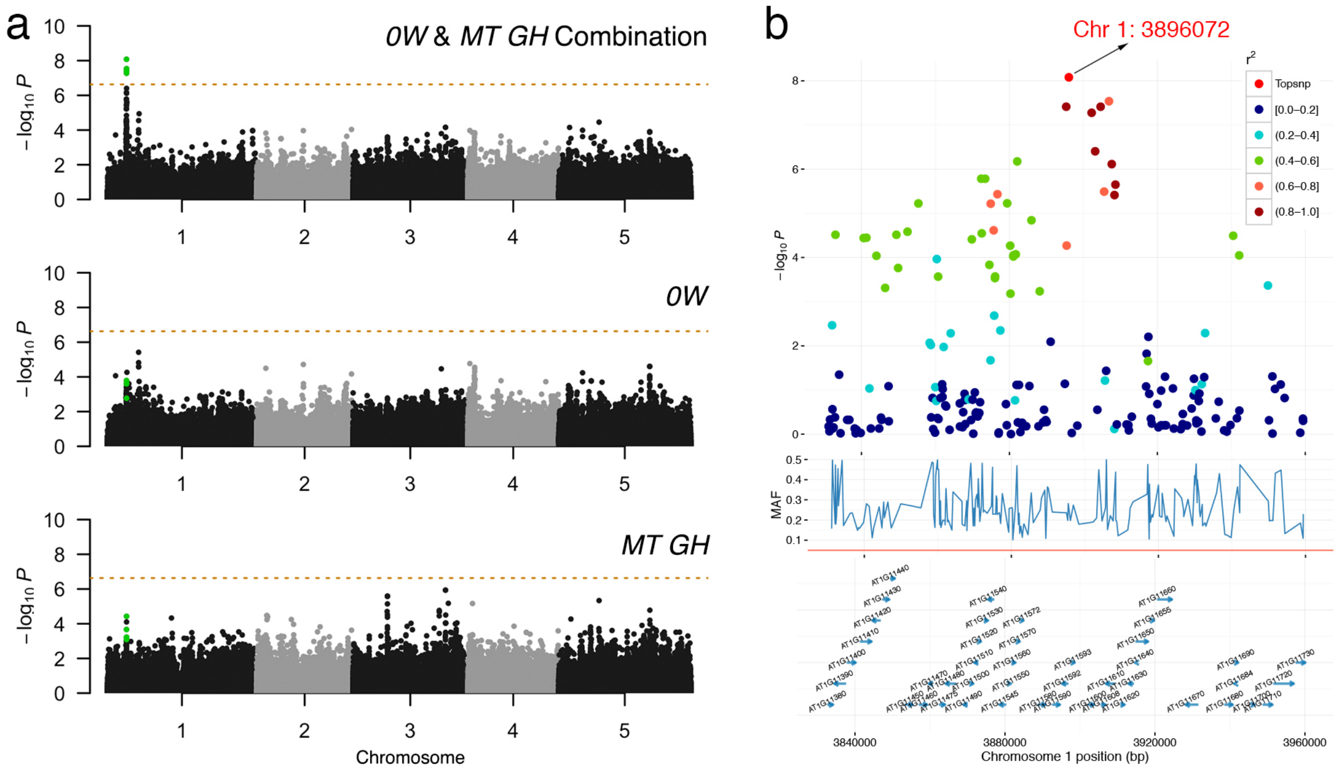


**Figure S3: Bivariate genome-wide association analysis of two developmental traits, 0W: Days to flowering time (FT) under Long Day (LD) without vernalization, MT GH: Maturation period.** (a) Manhattan plots comparison of bivariate and univariate analysis results, where the novel variants are only discoverable when combining two phenotypes are shown in green. The horizontal dashed line represents a 5% Bonferroni-corrected genome-wide significant threshold. (b) Zooming in on the novel locus detected using bivariate analysis. $r$: linkage disequilibrium measured as the correlation coefficient between the top variant and each variant in the region.

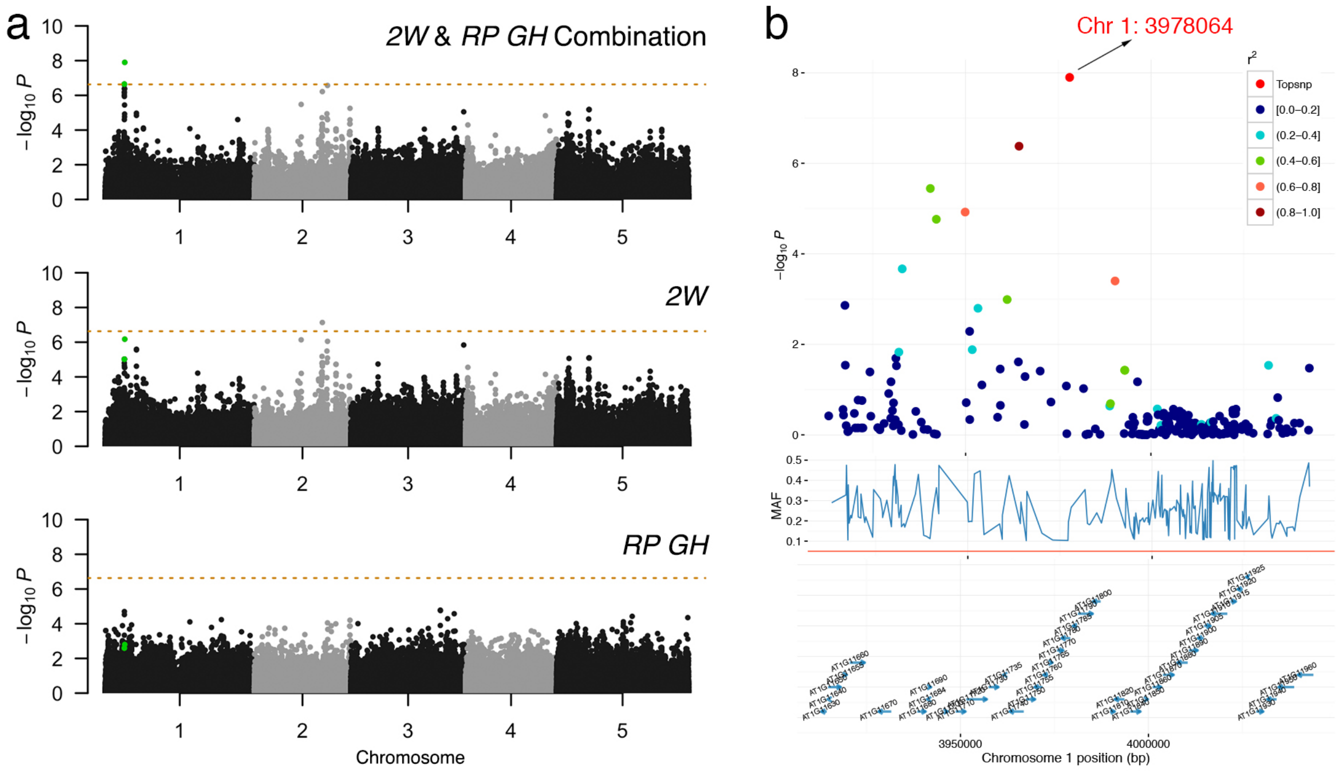


**Figure S4: Bivariate genome-wide association analysis of two developmental traits, 2W: Days to flowering time (FT) under long day (LD) with vernalized for 2 wks at 5°C, 8 hrs daylight, RP GH: Reproduction period.** (a) Manhattan plots comparison of bivariate and univariate analysis results, where the novel variants are only discoverable when combining two phenotypes are shown in green. The horizontal dashed line represents a 5% Bonferroni-corrected genome-wide significant threshold. (b) Zooming in on the novel locus detected using bivariate analysis. $r$: linkage disequilibrium measured as the correlation coefficient between the top variant and each variant in the region.

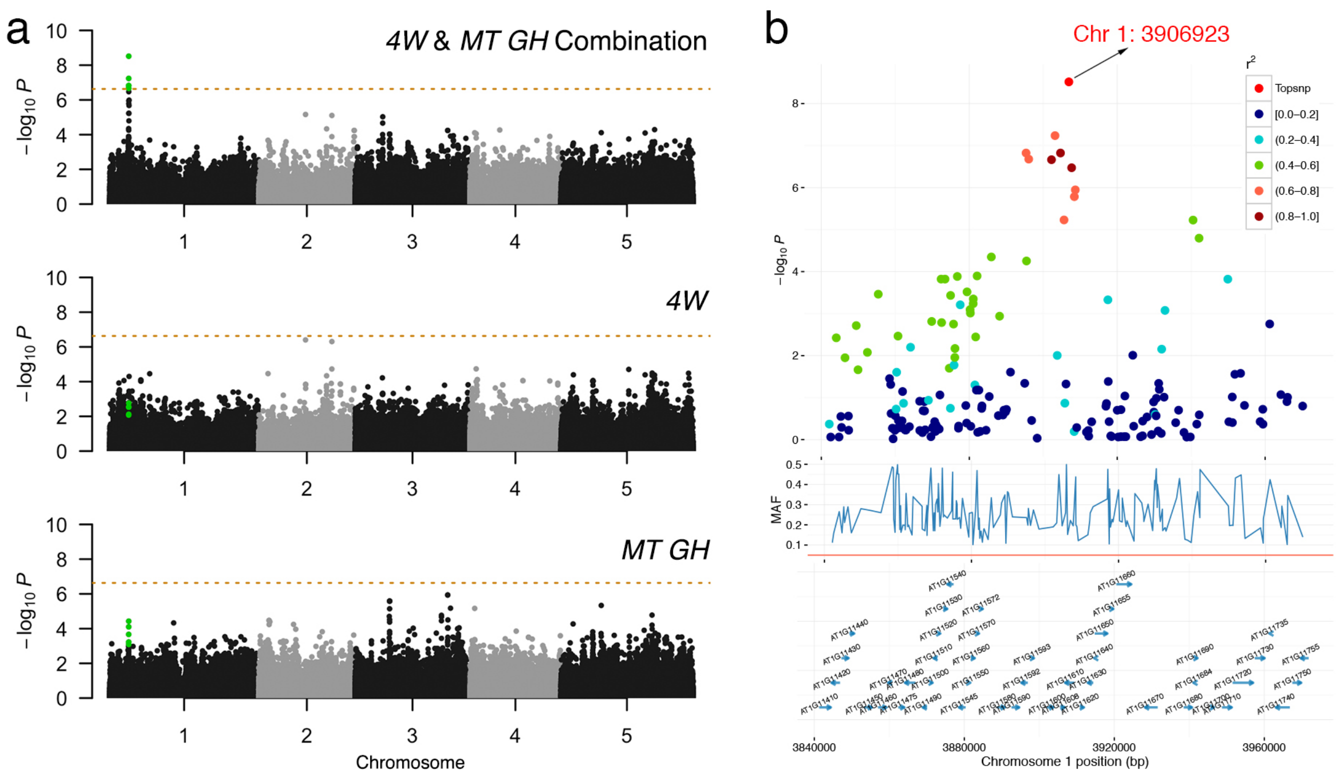


**Figure S5: Bivariate genome-wide association analysis of two developmental traits, 4W: Days to flowering time (FT) under long day (LD) with vernalized for 4 wks at 5°C, 8hrs daylight, MT GH: Maturation period.** (a) Manhattan plots comparison of bivariate and univariate analysis results, where the novel variants are only discoverable when combining two phenotypes are shown in green. The horizontal dashed line represents a 5% Bonferroni-corrected genome-wide significant threshold. (b) Zooming in on the novel locus detected using bivariate analysis. $r$: linkage disequilibrium measured as the correlation coefficient between the top variant and each variant in the region.

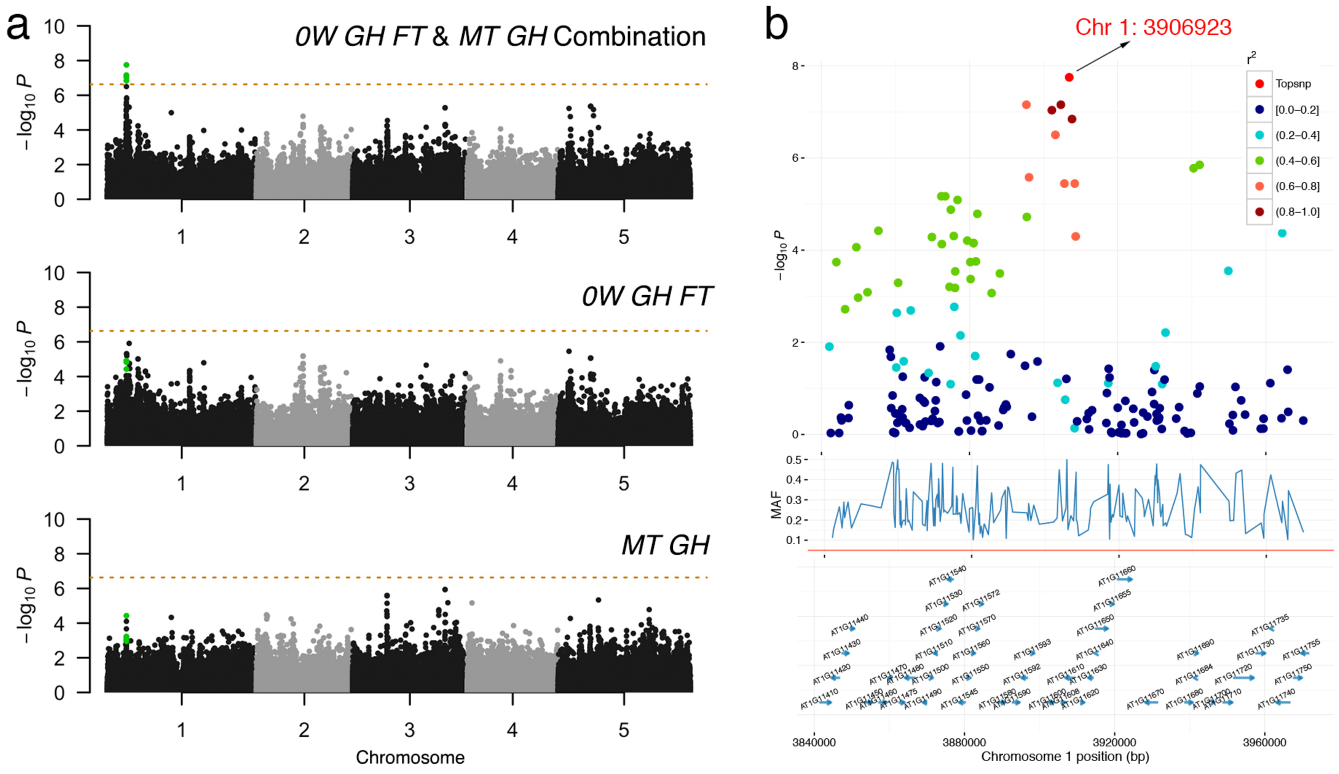


**Figure S6: Bivariate genome-wide association analysis of two developmental traits, 0W GH FT: Days to flowering time (FT), MT GH: Maturation period.** (a) Manhattan plots comparison of bivariate and univariate analysis results, where the novel variants are only discoverable when combining two phenotypes are shown in green. The horizontal dashed line represents a 5% Bonferroni-corrected genome-wide significant threshold. (b) Zooming in on the novel locus detected using bivariate analysis. $r$: linkage disequilibrium measured as the correlation coefficient between the top variant and each variant in the region.

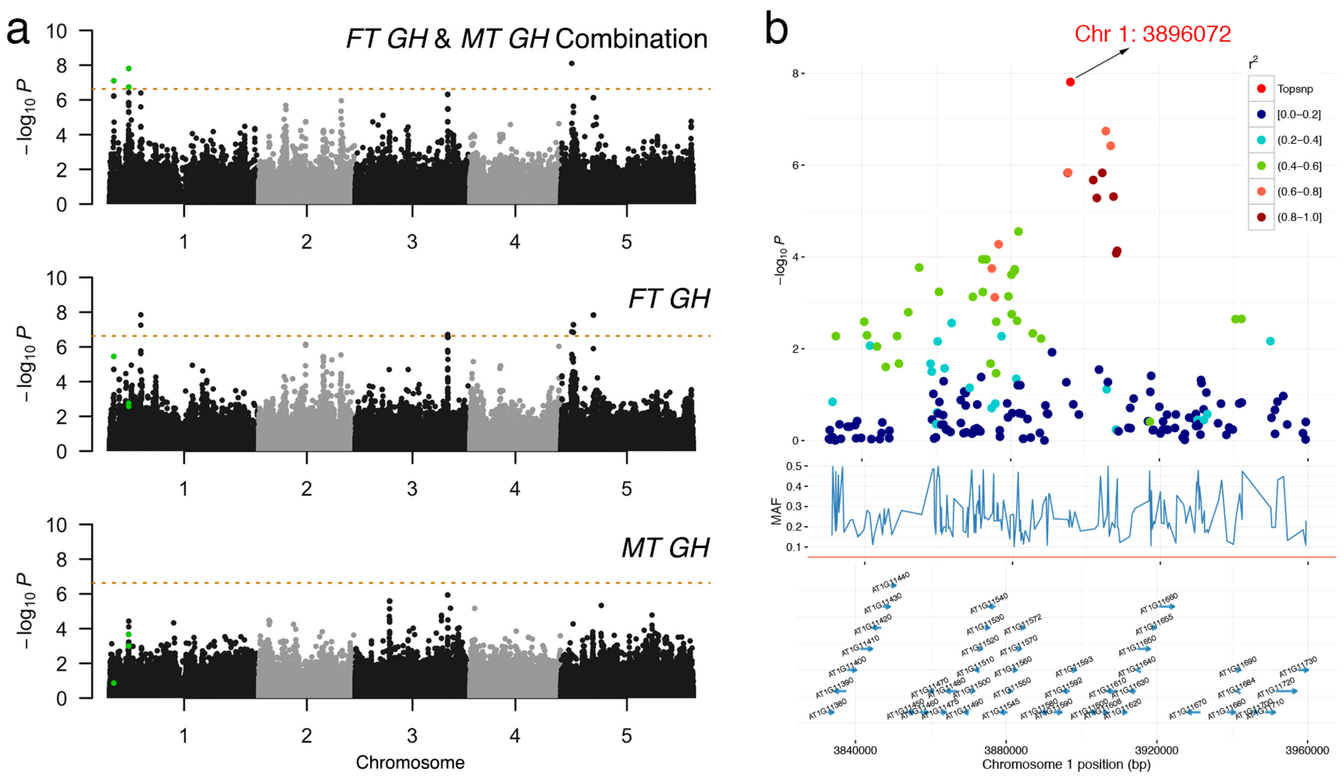


**Figure S7: Bivariate genome-wide association analysis of two developmental traits, FT GH: Days to flowering (greenhouse), MT GH: Maturation period.** (a) Manhattan plots comparison of bivariate and univariate analysis results, where the novel variants are only discoverable when combining two phenotypes are shown in green. The horizontal dashed line represents a 5% Bonferroni-corrected genome-wide significant threshold. (b) Zooming in on the novel locus detected using bivariate analysis. $r$: linkage disequilibrium measured as the correlation coefficient between the top variant and each variant in the region.

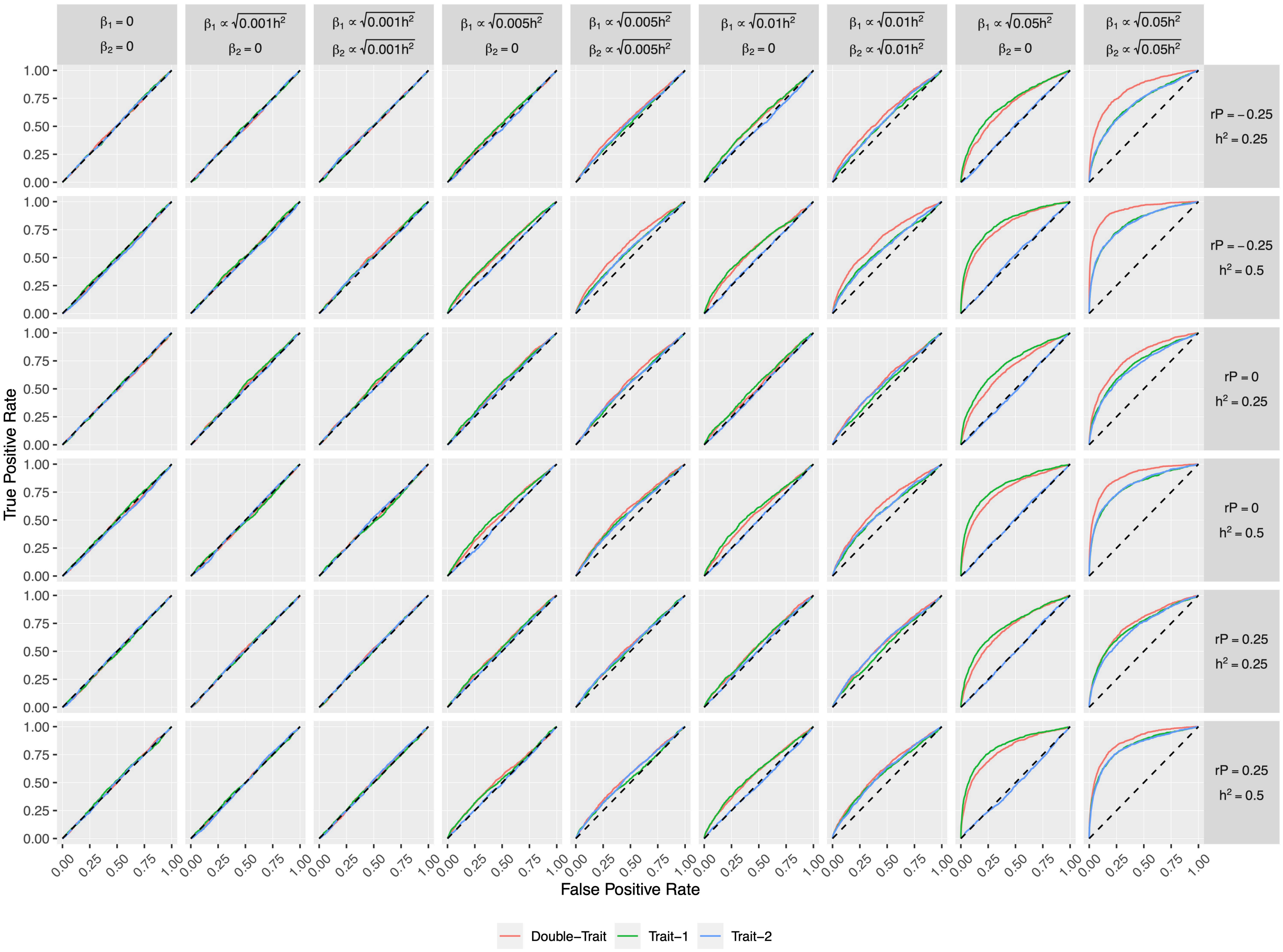


**Figure S8: Comparisons of single-trait and double-trait GWAS results of the target SNP under different simulated scenarios using 199 natural *Arabidopsis thaliana* inbred lines genotype data.** For each scenario, ROC curves display the true positive rate against the false positive rate with single- and double-trait GWAS methods from 1000 simulations. The $\beta$ represents the effect size of the target SNP in each phenotype. The heritability ($h^2$) of each phenotype was set to 0.25 and 0.5, based on 10% genome-wide SNPs randomly to simulate population structure. The phenotypic correlation ($r_P$) was set to -0.25, 0, and 0.25, respectively.

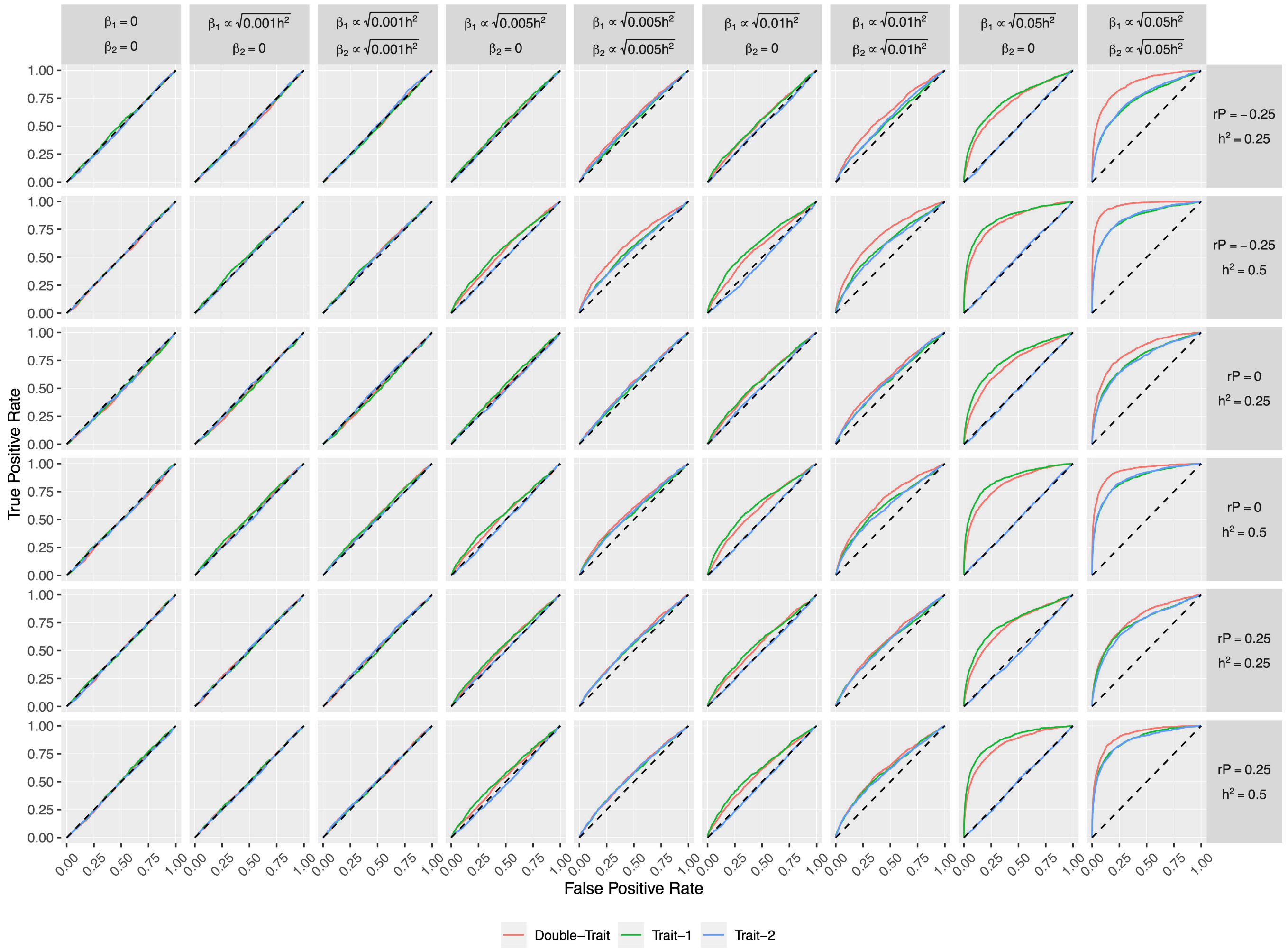


**Figure S9: Comparisons of single-trait and double-trait GWAS results of the target SNP under different simulated scenarios using 199 natural *Arabidopsis thaliana* inbred lines genotype data with half of the population structure.** For each scenario, ROC curves display the true positive rate against the false positive rate with single- and double-trait GWAS methods from 1000 simulations. The $\beta$ represents the effect size of the target SNP in each phenotype. The heritability ($h^2$) of each phenotype was set to 0.25 and 0.5, based on 10% genome-wide SNPs randomly to simulate population structure. The phenotypic correlation ($r_P$) was set to -0.25, 0, and 0.25, respectively.

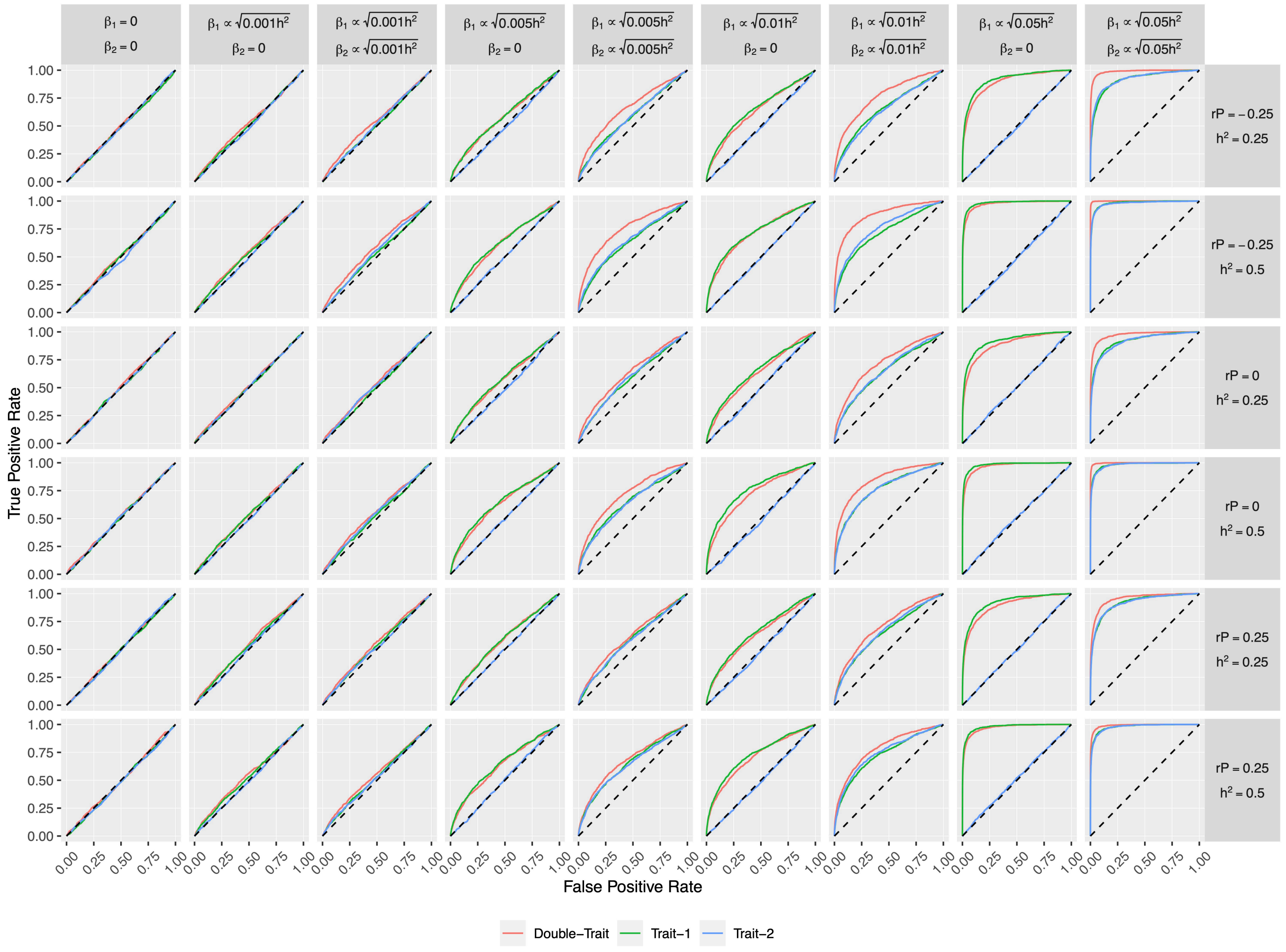


**Figure S10: Comparisons of single-trait and double-trait GWAS results of the target SNP under different simulated scenarios using genotype data of the 1001 genomes project for *Arabidopsis thaliana*.** For each scenario, ROC curves display the true positive rate against the false positive rate with single- and double-trait GWAS methods from 1000 simulations. The $\beta$ represents the effect size of the target SNP in each phenotype. The heritability ($h^2$) of each phenotype was set to 0.25 and 0.5, based on 10% genome-wide SNPs randomly to simulate population structure. The phenotypic correlation ($r_P$) was set to -0.25, 0, and 0.25, respectively.

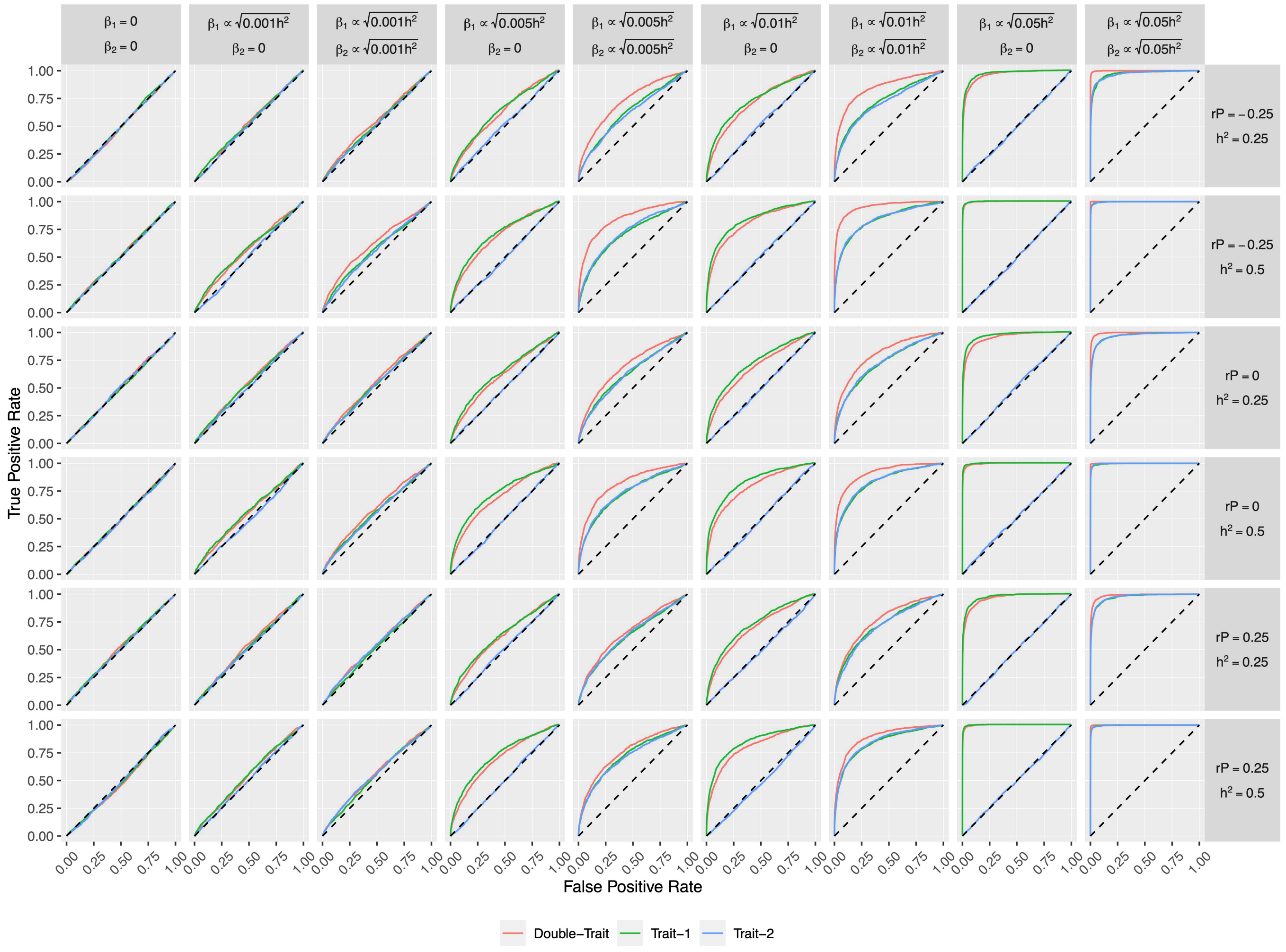


**Figure S11: Comparisons of single-trait and double-trait GWAS results of the target SNP under different simulated scenarios using genotype data of the 1001 genomes project for *Arabidopsis thaliana* with half of the population structure.** For each scenario, ROC curves display the true positive rate against the false positive rate with single- and double-trait GWAS methods from 1000 simulations. The $\beta$ represents the effect size of the target SNP in each phenotype. The heritability ($h^2$) of each phenotype was set to 0.25 and 0.5, based on 10% genome-wide SNPs randomly to simulate population structure. The phenotypic correlation ($r_P$) was set to -0.25, 0, and 0.25, respectively.

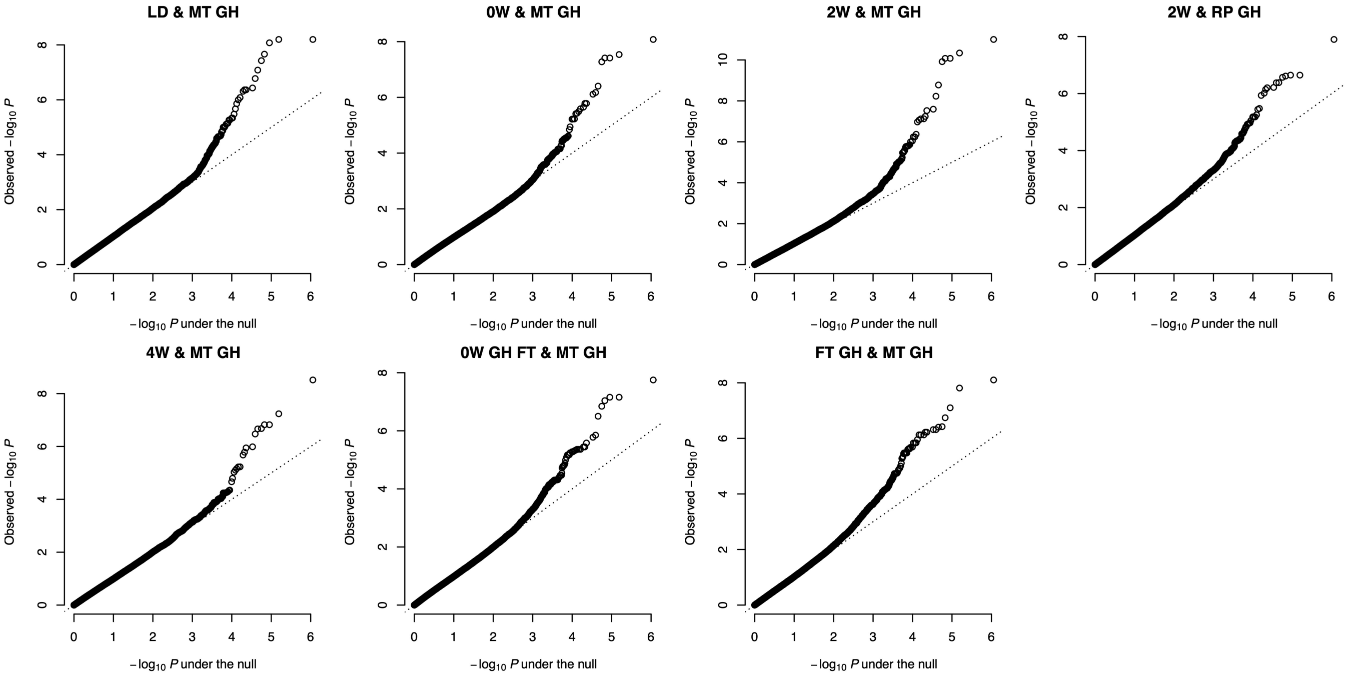


**Figure S12: Quantile-quantile (Q-Q) plots showing the $-log_{10}P$ under the null on the X-axis and the observed $-log_{10}P$ on the Y-axis.** The trait combinations are indicated at the top of each plot. Detailed information about these traits can be found in Table S1.

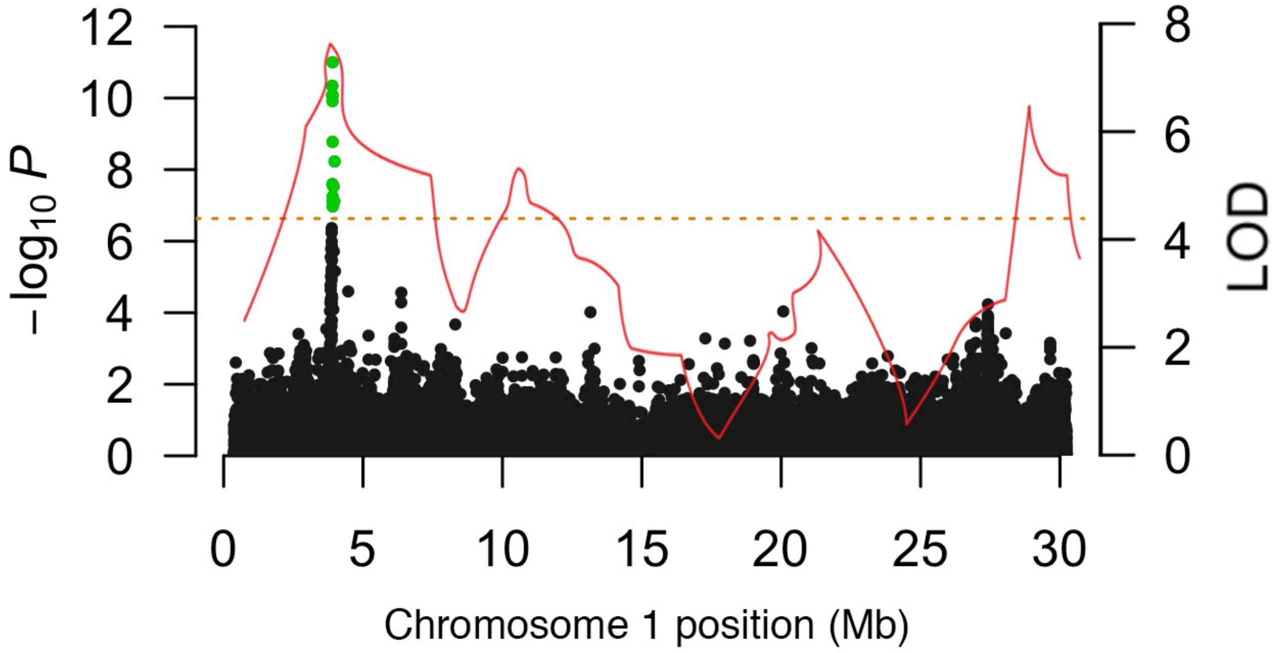


**Figure S13: Overlapping between QTL mapping and double-trait GWAS result.** The curve shows stepwise LOD profiles in chromosome 1 generated from a QTL mapping study using a cross between Italy and Sweden population analyzed by Dittmar et al. (2014) (reproduced by depicting the curvature of Figure 3a therein). The Manhattan plot shows the chromosome 1 signal in our bivariate analysis.
